# Identification of new interactors of eIF3f by endogenous proximity-dependent biotin labelling in human muscle cells

**DOI:** 10.1101/2025.02.10.636994

**Authors:** Lionel Tintignac, Nitish Mittal, Shahidul Alam, Meric Ataman, Yusuf I. Ertuna, Thomas Bock, Beat Erne, Mihaela Zavolan, Michael Sinnreich

**Affiliations:** Departments of Neurology and Biomedicine, Pharmazentrum, University and University Hospital of Basel, Klingelbergstrasse 50, 4056 Basel, Switzerland; Biozentrum, University of Basel, Spitalstrasse 41, 4056 Basel, Switzerland; Centre for Organismal Studies (COS), Heidelberg University, Heidelberg, Germany; Swiss Institute of Bioinformatics, Lausanne, Switzerland

## Abstract

Regulation of protein synthesis is central to maintaining skeletal muscle integrity and its understanding is important for the treatment of muscular and neuromuscular pathologies. The eIF3f subunit of the translation initiation factor eIF3 has a key role, as it stands at the crossroad between protein-synthesis-associated hypertrophy and MAFbx/atrogin-1-dependent. To decipher the molecular mechanisms underpinning the role of eIF3f in regulating muscle mass, we established a cellular model that enables interrogation of eIF3f functionality via identification of proximal interactors. Using CRISPR-Cas9 molecular scissors, we generated single cell clones of immortalised human muscle cells expressing eIF3f fused to the BirA biotin ligase (eIF3f-BioID1 chimera) from the endogenous *EIF3F* locus. Biotinylated proteins, representing interactors of eIF3f in nanometer range distance, were identified by streptavidin pull-downs and mass spectrometry. In both proliferating and differentiated muscle cells, the eIF3f-BioID1 chimera co-sedimented with ribosomal complexes in polysome profiles and interacted mainly with components of the eIF3 complex, and with the eIF4E, eIF4G, and eIF5 initiation factors. Surprisingly, we identified several nucleus-localised interactors of eIF3f, and the immunofluorescence analyses revealed a previously unknown nuclear localization of eIF3f in both myoblasts and myotubes. We also identified novel cytoplasmic partners of eIF3f, responsible for the maintenance of skeletal muscle ultrastructure (sarcomeric/Z-disc (SYNPO2) bound proteins) and proteins of the lysosomal compartment (LAMP1). The established tagging system should be useful to further advance studies of eIF3f function in hypertrophic and atrophic conditions in skeletal muscle.

## INTRODUCTION

Protein synthesis is central to maintaining skeletal muscle integrity and its deregulation leads to muscle wasting, a hallmark of muscle pathologies such as sarcopenia and neuromuscular diseases^1^. At the molecular level, catabolic and anabolic signalling pathways have been shown to converge on sensor proteins such as the eukaryotic translation initiation factor eIF3f. This core component of eIF3 is rapidly degraded by the ubiquitin proteasome machinery during early responses to muscle insults, ultimately initiating muscle wasting^2^. Genetic ablation of eIF3f is embryonically lethal in mice and its reduced expression in heterozygous EIF3F+/- animals correlates with a reduced muscle mass^3^. Intriguingly, eIF3f downregulation is often observed in cancer cells where it was proposed to be a tumour suppressor candidate^4^. Such functional polymorphism made eIF3f an attractive candidate to study in the context of physiological changes in the muscle.

The multistep process of mRNA translation is divided into three phases: initiation, elongation and termination. The process is predominantly regulated at the initiation phase by various initiation factors such as eIF3, which also plays a role in elongation and termination^5,6^. The eIF3 complex is involved in both canonical (m^7^G-cap-dependent) and non-canonical translation initiation^7^. In canonical translation, the capped mRNA associates with the eIF4F complex of three proteins, eIF4E, eIF4A, and eIF4G, with eIF4E recognizing the m^7^G-cap. In parallel, eIF3 and other initiation factors, including the ternary complex (TC) bearing the initiator Methionyl-tRNA (eIF2·GTP·Met-tRNAi), associate with the 40S ribosomal subunit to form the 43S pre-initiation complex (PIC). The 43S PIC then attaches to the 5’ end of mRNA bound by eIF4F, forming the 48S PIC that initiates scanning of the mRNA 5’-UTR for the AUG start codon. In non-canonical translation initiation, operating, for example, in many viruses, the 5’ UTR of mRNA contains internal ribosomal entry sites (IRES). These are recognised by the 40S ribosomal subunit and the eIF3 complex, and do not require the full complement of initiation factors for translation initiation. Recently, additional non-canonical initiation mechanisms have been described (reviewed in ^8^). Interestingly, one of them involves the eIF3 subunit eIF3d, which was shown to recognize the mRNA m^7^G-cap, with this activity being regulated by eIF3d phosphorylation during stresses ^9,10^.

eIF3 is a large complex (>800 kDa) composed in vertebrates of 13 subunits (a through m)^11^. Its octameric structural core is composed of two sub-complexes: the six subunit (a, c, e, l, k and m) PCI (proteasome, COP9/signalosome, eIF3) complex, and a dimeric MPN (Mpr1-Pad1-N-terminal) complex consisting of eIF3f and h. The 5-lobed shaped eIF3 core shares structural similarity with the 19S proteasome lid and the COP9 signalosome. Structural studies in HEK293T cells suggested that the assembly of the complex is nucleated around the largest subunit, eIF3a, by subunits b, g, and i, followed by the gradual assembly of the helical bundle (7 helices in total) involving C-terminal helical regions of eIF3f and other core subunits^12,13^. Four subunits (b, d, g and i) are stably attached to the core octamer, with subunits b and g containing the RNA Recognition Motives (RRMs) that help eIF3 complex bind to RNA. eIF3j is only loosely attached to the complex (reviewed in ^14^). Among the 13 subunits, eIF3f was shown to display specific and unique functions in skeletal muscle cells.

Unbalanced protein homeostasis (“proteostasis”) occurs in most, if not all, muscle pathologies, caused by dysregulation of protein synthesis, degradation, folding, and/or trafficking^15^. Atrogenes, a family of genes regulating the ubiquitin-proteasome pathway (UPP) and the autophagy pathway, are activators of muscle wasting in mice and humans. MAFbx, an F-box-containing member of the atrogene family, was found to target eIF3f for ubiquitin-dependent degradation during atrophy. The MAFbx knock-out muscle was found to spare ∼40% of size in different atrophy models^16^, while the overexpression of eIF3f in the muscle induced hypertrophy^2^. Since the execution of the atrophy program is accompanied by the expression of various components of the myocellular degradation machinery, we postulated that eIF3f may exert a myo-protective function. The reduction of eIF3f protein in heterozygous EIF3F^+/-^ mice leads to reduced mass of muscle, kidney, heart and brain^3^. Molecular characterization of EIF3F^+/-^ muscle showed disturbed growth signalling with reduced S6K1 and 4EBP1 phosphorylation levels demonstrating the reduction of mTORC1 activity^3^. Reduced polysomal and sub-polysomal fractions from quadriceps and gastrocnemius muscles of EIF3F^+/-^ mice demonstrated a global defect in protein synthesis. In humans, an autosomal recessive homozygous missense mutation of eIF3f (Phe232Val) was shown to result in 30% reduction in eIF3f protein expression and to be causative for neurodevelopmental disorders^17^. Reduction of eIF3f expression was also reported in melanocytic neoplasms, pancreatic cancers, breast and ovary cancers, and eIF3f together with eIF3e were the only members of the eIF3 complex found to be reduced in these cancers^18^. In Jurkat cells, siRNA induced depletion of eIF3f increases proliferation and protects cells from apoptosis whereas CMV-driven eIF3f overexpression is associated with rRNA degradation and ribosome decrease^19^. Together, these observations suggested that eIF3f is a tumor suppressor candidate. However, in colorectal cancer eIF3f is overexpressed and has a tumor-driving function that is independent of translation and involves a deubiquitinase activity encoded by the MPN/MOV34 motif, eIF3f shares with other members of the JAB1/MPN/MOV34 metalloenzyme (JAMM) family of deubiquitinases (DUBs)^20,21^.

Finally, a recent study in cancer cells investigating the effect of the individual eIF3 subunit knock-downs on translation (by polysome profiling) and protein output, demonstrated that despite marked functional differences on the eIF3 holo-complex, the eIF3a, b, e, and f subunits are essential for cancer cell proliferation and tumor growth^22^. In contrast to previous observations that eIF3f inhibits cancer cell protein synthesis via hnRNP K-eIF3f interaction^4^, this elegant study reported that only eIF3k depletion increases global translation through relieving repression of synthesis of ribosomal proteins, especially of RPS15A^22^.

To date our mechanistic knowledge of eIF3f stems from contradictory results obtained in cancer cells, where eIF3f expression has been associated with translation inhibition or activation^23^. Moreover, the eIF3f functions in human skeletal muscle are unknown which prompted us to establish a cellular model to interrogate eIF3f functionality based on the identification of its proximal interactors. To this end, we decided to employ the biotin ligase enzymatic strategy to tag eIF3f interactors *in cellulo*.

Proximity-dependent biotin identification (BioID) was developed to study protein-protein interaction (PPI) via the labelling of direct interactors of the protein of interest (POI) recorded during the labelling period. This technique has already been successfully used to identify PPI networks specific to subcellular organelles^24^, macrostructures^25^ and signalling pathways^26^. The bacterial BirA biotin ligase from *Escherichia coli* was optimised by modification of its catalytic site (denoted as BirA*; mutant R118G)^24^ for *in situ* proximity-dependent labelling assay. The technique has already been applied to the eIF3a protein, as a virally expressed form of eIF3a- BirA* was used to identify the interactome in colon cancer cells^27^.

Here, we first designed and validated the activity of eIF3f-BioID chimera after transient expression in HEK293 cells in the presence of exogenous biotin. We identified all components of the PCI complex (except eIF3k) and the MPN, demonstrating that C-terminal addition of BirA* does not interfere with the assembly of a functional eIF3 complex. To alleviate the caveat of CMV or viral-driven overexpression of the chimera, we took advantage of the CRISPR-Cas9 system to create a locus-specific EIF3F-BioID fusion in human muscle cells. We then characterised the muscle-specific, endogenous, proximal interactome of eIF3f in both proliferating and differentiated human muscle cells, in physiological conditions. We identified the eIF3 core octamer components as major interactors of eIF3f and demonstrated the association of eIF3f with both monosome and polysome fractions. Among the eIF3f interactors we also identified numerous nuclear proteins along with proteins involved in maintenance of sarcomeric/cytoplasmic structure, indicating that eIF3f is present in both nuclear and cytoplasmic compartments. While in mononucleated muscle cells the protein was distributed diffusely, in multinucleated cells, its distribution pattern was more granular. The set of eIF3f was larger in differentiated compared to proliferating cells and included the lysosomal protein LAMP1. Our work reveals an unanticipated pattern of eIF3f distribution in muscle cells and suggests that eIF3f may regulate localised protein synthesis.

## MATERIALS AND METHODS

### Cell culture and transfection

HEK293 cells were cultured in DMEM glutamax high glucose (4.5g/L; Gibco 61965-026) supplemented with 10% FCS (Gibco A5256701) and Penicillin-Streptomycin (Sigma-Aldrich P4333). Transfections were achieved with JetPEI (PolyPlus 101-10N) following manufacturer’s recommendation. U16 immortalised (CDK4-hTERT) human cells (generous gift from Wright WE laboratory (ref. ^28^)) were cultured on 0.01% gelatin dishes. Myoblasts (MB) were maintained in proliferation in 4:1 ratio of DMEM glutamax/medium 199 (Gibco 22340-020), 20mM Hepes (MIMED 5-31F00H), 0.03ug/mL Zinc Sulfate, 121.4ug/mL Vitamin B, 0.055ug/mL Dexamethasone, 2.5ng/mL HGF (Preprotech 100-39H), 10ng/mL human bFGF (Preprotech 100-18B) supplemented with Penicillin-Streptomycin (Sigma-Aldrich P4333). Myotubes (MT) were obtained after induction of differentiation of 80% confluent myoblasts with 3 washes in 1X PBS and 6 days of culture in 4:1 DMEM glutamax/medium 199 supplemented with 10mg/L Insulin (Sigma I9278), 0.02M Sodium Pyruvate (Sigma 58636) and 2% Horse serum (Gibco 16050-130). MB cells were transfected by electroporation with the Neon^TM^ (Thermo Fisher Scientific) system following manufacturer recommendations.

### Cloning

All plasmid backbones mentioned thereafter are listed in table 1.

**Table 1.** Plasmids used in the study.

| Plasmid backbone | Source |
| --- | --- |
| pcDNA3.1 mycBioID | Addgene, # 35700 |
| pcDNA3.1 destination MCS-BirA(R118G)-HA | Addgene, # 53581 |
| pcDNA 3.1 MCS-BioID2-HA | Addgene, # 74224 |
| pcDNA 3.1 myc-BioID2-MCS | Addgene, # 74223 |
| pSpCas9(BB)-2A-GFP (PX458) | Addgene, # 48138 |
| pSpCas9(BB)-2A-Puro (PX459) | Addgene, # 62988 |
| pU6-(BbsI)_CBh-Cas9-T2A-BFP-P2A-Ad4E1B | Addgene, # 64218 |
| pUC19 | Addgene, # 50005 |

PCR amplified human EIF3F (ref. ^2^) was cloned into the pDONOR/Zeo plasmid of the gateway cloning system (Invitrogen) which was subsequently used to subclone EIF3F into the pDEST- RK5-Myc vector. pZeo-EIF3F was used to generate eIF3f–BioID with both BirA(R118G)-HA destination and pcDNA3.1 mycBioID vectors. Stop-less hEIF3F was PCR amplified and ligated in Age1/BamH1 double digested MCS-BioID2-HA and ATG-less hEIF3F was PCR amplified and ligated in XhoI/HindIII of Myc-BioID2-MCS. All plasmids were amplified in DH5α competent cells, purified using ZymoPure plasmid kit (Zymo research) and DNA sequence were confirmed by sequencing.

### Establishment of eIF3f-BioID1 expressing myoblasts

All primers and plasmid backbones mentioned thereafter are listed in tables 1 and 2. Benchling software (retrieved from https://benchling.com) was used to design oligonucleotides for expression of two different sgRNAs (sg1 and sg2) targeting upstream and downstream sequences of the Ex10 of human gene *EIF3F* (ENST00000533626.5) locus at chromosome 11p15.4. After annealing, the sgRNAs were subcloned in pSpCas9(BB)-2A-GFP. Plasmids were amplified, purified (Nucleobond) and sequenced, and selected clones were transiently transfected in HEK293. 48hrs after transfection, GFP positive cells were enriched by fluorescence activated cell sorting (FACS). Extracted genomic DNA (gDNA) (Qiagen) was genotyped by gel electrophoresis separation of the PCR product (NEBNext High-Fidelity PCR master Mix) amplified with EIF3F screening primer pair (EIF3F_scrn_F and EIF3F_scrn_R, see table 2). Additional assays were carried out to visualise the CRISPR induced mismatch of EIF3F Ex10 surrounding region. A pSpCas9(BB)-2A-Puro plasmid, where puromycin resistance gene was substituted by the cherry fluorescent protein gene, was used to subclone one of the sgRNAs before co-transfection in HEK293 cells as described above. Double positive cells (DP; GFP and mCherry) gDNA were used for survivor assay and sgRNAs (sg1 and sg2) were selected. Gibson assembly system (NEB) was used to assemble the donor DNA between HindIII and KpnI restriction site of pUC.19 backbone, composed of 6 PCR amplified fragments (supplementary figure 3) encoding left (LHA) and right (RHA) homology arms flanking the EIF3F Ex10 in frame with BioID-HA-T2A-BFP-stop for which the T2A-BFP was subcloned from pU6-(BbsI)_CBh-Cas9-T2A-BFP-P2A-Ad4E1B plasmid. The generated construct was electroporated into U16 muscle cells with the Neon electroporation system (Invitrogen). Cells were treated with 1μM Scr7 (Selleckchem) 6hrs after transfection to inhibit non-homologous DNA end joining. Double positive cells expressing GFP and mCherry were selected using FACS before gDNA extraction and genotyping. Finally, single BFP+ cells were FACS isolated and propagated. Single cell clones were validated using TOPO cloning of EIF3F Ex10 PCR product and sequencing.

**Table 2.**
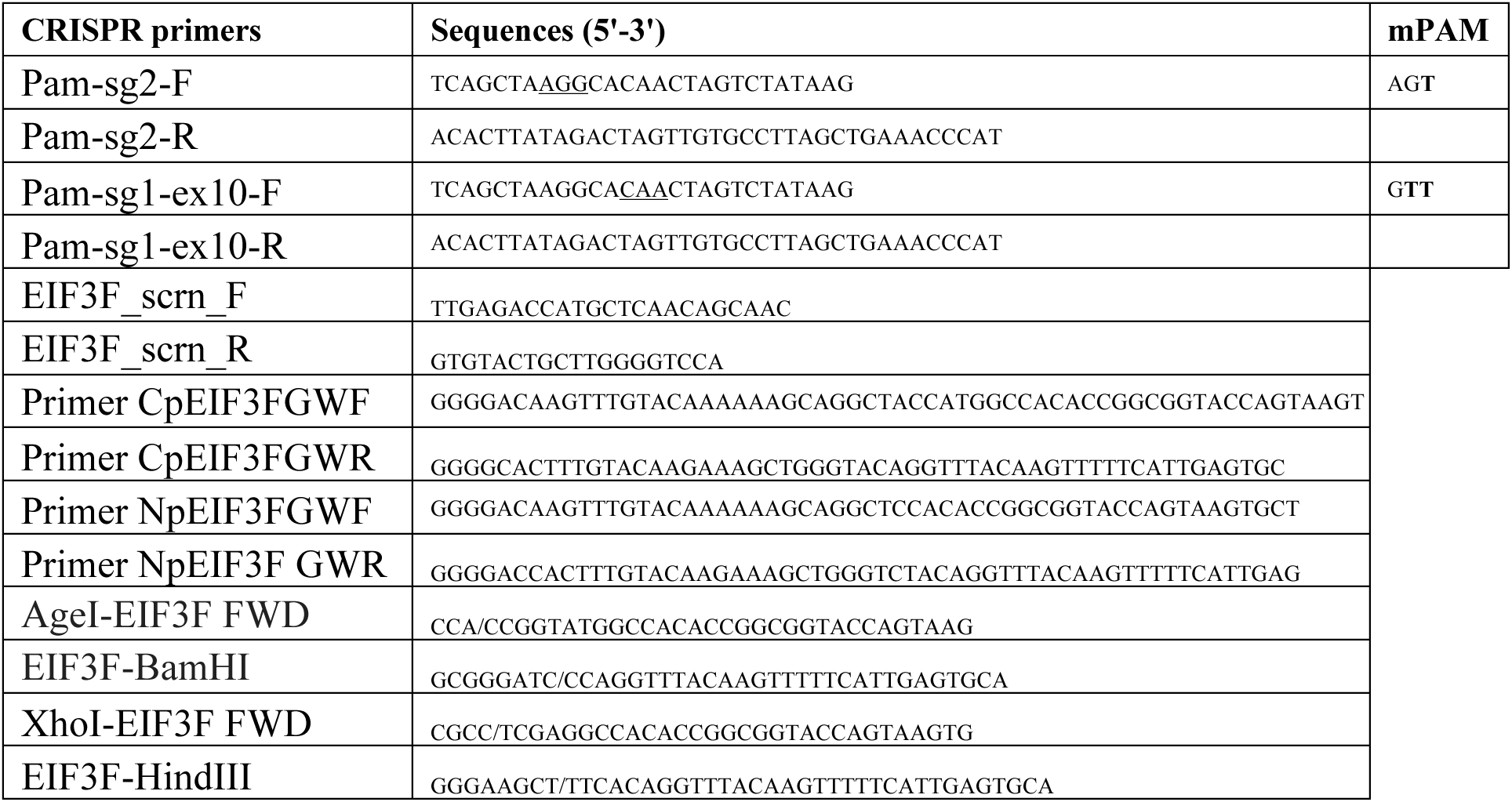
Primers used in the study.

### Cell lysis and Immunoprecipitation

Whole cell lysis was achieved with 10% glycerol, 50mM Tris-HCl pH7.5, 150nM NaCl, 1% triton, 1mM EDTA, 0.2mM Na3VO4, proteases inhibitor cocktail (cOmplete Mini EDTA free (Roche 11836170001) and PhoSTOP (Roche 04906837001). Upon short sonication at 4°C, the lysate was cleared by centrifugation at 10’000rpm for 10min (minute) at 4°C and protein concentration measured with Pierce BCA 23225 assay (Thermo Scientific) on a Tecan Infinite reader. For immunoprecipitation, equal amounts of protein (500μg) were incubated O/N on a rotating wheel with primary or irrelevant control antibody in 500μL of the same buffer. 20μL of 50/50 v/v mixture of Dynabeads coupled Protein G (Invitrogen Ref10004D)/Protein A (Pierce Ref 88846) were added for 1hr (hour) on a rotating wheel at 4°C. Beads were collected with a magnetic separation rack and washed 4 times with lysis buffer before addition of 25μL of 1X Laemmli buffer, boiling for 10min at 98°C before separation on SDS-PAGE gel.

### Western blotting

Protein samples were denatured upon addition of 5X Laemmli buffer and boiling for 10min at 98°C before separation on SDS-PAGE gel/gradient gel (Mini Protean 4-20% BioRAD #4561093). Proteins were later transferred (liquid) on nitro-cellulose membranes (Amersham Protran 0.45µm, Cytivia 10600002). Unspecific blocking was achieved by 1 hr incubation of the membrane at room temp in 5% bovine serum albumin (BSA Sigma A9647)/0.1% Tween (Tween20 Sigma P7949) in Tris-glycine buffer. Upon incubation with the primary antibody and washes, secondary antibody coupled to horseradish peroxidase were visualised with ECL chemiluminescence reagent on Fusion FX (Vilber) machine. Secondary antibodies coupled to infrared dyes were detected using Odyssey CLx (Li-COR), analysed and quantified with Image Studio 4.0 software. All antibodies used are listed in table 3.

**Table 3.** Antibodies used in the study.

| Antibody | Reference | Supplier |
| --- | --- | --- |
| anti-eIF3a | PA5-17212 | Invitrogen |
| anti-eIF3f | A303-005A | BETHYL |
| anti-eIF3f | ab176853 | Abcam |
| anti-eIF3l | PA5-99288 | Invitrogen |
| anti-eIF4E (C46AH6) | #2067T | CST |
| anti-eIF5 (D5G9) | #13894 | CST |
| anti-rps14 | 16683-1-AP | Proteintech |
| anti-Lamin B1 | PA5-19468 | Invitrogen |
| anti-LAMP1_D4O1S | #15665 | CST |
| anti-LAMP1_D2D11 | #9091 | CST |
| anti-Alpha-Actinin | A7732 | Sigma |
| anti-puromycin 12D10-AF647 | MABE343-AF647 | Sigma Aldrich |
| Rabbit anti-mouse IgG-HRP | ab6728 | abcam |
| Goat anti-rabbit IgG-HRP | ab6721 | abcam |
| IRDye® 800CW Goat anti-Mouse IgG | 926-32210 | LiCORbio |
| IRDye® 800CW Goat anti-Rabbit IgG | 926-32211 | LiCORbio |
| IRDye® 680RD Goat anti-Mouse IgG | 926-6807 | LiCORbio |
| IRDye® 680RD Goat anti-Rabbit IgG | 926-68071 | LiCORbio |
| Rabbit IgG isotype control | #31235 | Invitrogen |
| MouseIgG1 isotype Control | MA5-14453 | Invitrogen |
| Goat anti-rabbit 488 | 111-545-144 | Jackson Imm |
| Goat anti-mouse 488 | 111-545-146 | Jackson Imm |
| Goat anti-mouse 568 | AZ1124 | LiseTech |
| Donkey anti-rabbit Cy5 | 711-175-152 | Jackson Imm |

### Cell fractionation

The protocol was adapted from Dimauro et al^29^. Briefly, MB or MT cells were homogenised at 4°C using a Teflon pestle in 450μl of STM buffer (250mM sucrose, 50mM Tris–HCl pH7.4, 5mM MgCl2, protease and phosphatase inhibitor cocktails). Upon differential centrifugation, the pellet containing nuclei was lysed in NET buffer (20mM HEPES pH 7.9, 1.5mM MgCl2, 0.5M NaCl, 0.2mM EDTA, 20% glycerol, 1% TritonX-100, protease and phosphatase inhibitors), sonicated at high setting (80%) before centrifugation at 9,000g for 30min at 4°C and collection of the supernatant as the “nuclear fraction”. Cytosol and microsomes found in the supernatant of the first differential centrifugation were precipitated in cold 100% acetone at −20°C and collected as a pellet resuspended in STM buffer and 1X Laemmli buffer.

### Affinity Pulldown Sample Preparation

#### Collection and purification of biotinylated proteins

10cm dishes of cultured cells were lysed with 500uL of BioID buffer (50 mM Tris, pH 7.4, 500 mM NaCl, 0.4% SDS, 5 mM EDTA, 1 mM DTT, 1X complete protease inhibitor (Roche 1xPhoStop). Following protocol initially published by Roux et al., cells were sonicated (5 up and down) and triton X-100 was added to 2% final concentration and diluted by addition of 1 v/v of 50mM Tris pH7.5. Lysates were cleared by centrifugation at 16,000g for 10min at 4°C. Supernatants were distributed to a new Eppendorf tube containing 30uL of 50% (v/v) Dynabeads MyOne Streptavidin C1 (Invitrogen #65001) previously washed twice with BioID buffer. After O/N incubation on a rotating wheel at 4°C the beads were collected with a magnetic rack and washed twice with wash buffer 1 (2% SDS qsp H2O); once with washed buffer 2 (0.1% deoxycholate, 1% Triton X-100, 500mM NaCl, 1mM EDTA, 50mM Hepes, pH 7.5); once washed with buffer 3 (250mM LiCl, 0.5% NP-40, 0.5% deoxycholate, 1mM EDTA, 10mM Tris, pH 8.1); twice washed with buffer 4 (50mM Tris pH 7.4, 50mM NaCl) and finally washed twice in 50mM NH4HCO3. Biotinylated pulled down proteins were released on a magnetic rack in 40uL of 5%SDS and 10mM biotin after heating 10min at 98°C. Samples were stored at -80°C for later MS/MS analysis.

### Mass spectrometry protein analysis

Purified biotinylated samples (1% Sodium deoxycholate (SDC), 10mM TCEP, 100mM Tris, pH8.5) were sonicated twenty cycles (30s on, 30s off per cycle) on a Bioruptor (Dianode). Following sonication, pulldown samples were reduced by TCEP at 95°C for 10min. Proteins were alkylated using 15mM iodoacetamide at 25°C for 30min and further digested using sequencing-grade modified trypsin (1/25 w/w, ratio trypsin/protein; Promega, USA) on a STRAP column (protifi.com) at 47° C for 1hr according to the manufacturer’s instructions. STRAP eluate containing peptide digests were dried under vacuum and peptides were stored at -20°C. Prior to use peptides were dissolved in 0.1% aqueous formic acid solution at a concentration of 0.2mg/ml.

For each sample, aliquots of 0.2μg of total peptides were subjected to LC-MS analysis using a Q Exactive Plus Mass Spectrometer fitted with EASY-nLC 1000 (both Thermo Fisher Scientific). Peptides were resolved using an EasySpray RP-HPLC column (75μm × 25cm, Thermo Fisher Scientific) at a flow rate of 0.2μL/min. The following gradient was used for peptide separation: from 5% B to 10% B over 5min to 35% B over 40min to 50% B over 10min to 95% B over 2min followed by 18min at 95% B. Buffer A was 0.1% formic acid in water and buffer B was 80% acetonitrile, 0.1% formic acid in water.

For data dependent acquisition (DDA) analysis each MS1 scan was followed by high-collision-dissociation (HCD) of the 10 most abundant precursor ions with dynamic exclusion for 45 seconds. For MS1, 3x10^6^ ions were accumulated in the Orbitrap cell over a maximum time of 100ms and scanned at a resolution of 70,000 FWHM (at 200m/z). MS2 scans were acquired at a target setting of 1x10^5^ ions, an accumulation time of 100ms and a resolution of 35,000 FWHM (at 200m/z). Singly charged ions and ions with unassigned charge state were excluded from triggering MS2 events. The normalised collision energy was set to 28%, the mass isolation window was set to 1.4 m/z and one microscan was acquired for each spectrum.

### Protein Identification and Label-free Quantification

The acquired raw-files were imported into the Progenesis QI software (v2.0, Nonlinear Dynamics Limited), which was used to extract peptide precursor ion intensities across all samples applying the default parameters. The generated mgf-files were searched using MASCOT with a target-decoy search strategy against a database containing normal and reverse sequences of the *Homo sapiens* proteome (UniProt, release date: 17.04.2020) and commonly observed contaminants (41484 protein entries) generated using the SequenceReverser tool from the MaxQuant software (Version 1.0.13.13). The following search criteria were used: full tryptic specificity was required (cleavage after lysine or arginine residues, unless followed by proline); 3 missed cleavages were allowed; carbamidomethylation © was set as fixed modification; oxidation (M) and lysine biotinylation (226 Da) were applied as variable modifications; mass tolerance of 10 ppm (precursor) and 0.02 Da (fragments). The database search results were filtered using the ion score to set the false discovery rate (FDR) to 1% on the peptide and protein level, respectively, based on the number of reverse protein sequence hits in the datasets.

Quantitative analysis results from label-free quantification were normalised and statically analysed using the SafeQuant R package v.2.3.4 ((https://github.com/eahrne/SafeQuant/) ref.^30^) to obtain protein relative abundances. This analysis included quantification of the total peak/reporter areas across all LC-MS runs, the summation of peak areas per protein and LC-MS/MS run, followed by calculation of protein abundance ratios. Only isoform-specific peptide ion signals were considered for quantification.

For “core eIF3f-eIF3” interactor identification, instead of global normalization a normalization against the protein abundances of the following four protein subunits was chosen: EIF3A (Q14152; EIF3CL(B5ME19); EIF3E (P60228); EIF3F (O00303) and EIF3H (O15372). The summarised protein expression values were used for statistical testing of differential protein expression between conditions. Here, empirical Bayes moderated t-tests were applied, as implemented in the R/Bioconductor limma package (http://bioconductor.org/packages/release/bioc/html/limma.html). The resulting per protein and condition comparison p-values were adjusted for multiple testing using the Benjamini-Hochberg method.

All LC-MS analysis runs were acquired from independent biological samples. To meet additional assumptions (normality and homoscedasticity) underlying the use of linear regression models and Student’s t-test, MS-intensity signals are transformed from the linear to the log-scale.

Unless stated otherwise linear regression was performed using the ordinary least square (OLS) method as implemented in *base* package of R v.3.1.2 (http://www.R-project.org/).

The sample size of three biological replicates was chosen assuming a within-group MS-signal Coefficient of Variation of 10%. When applying a two-sample, two-sided Student t-test this gives adequate power (80%) to detect protein abundance fold changes higher than 1.65 per statistical test. Note that the statistical package used to assess protein abundance changes, SafeQuant, employs a moderated t-test, which has been shown to provide higher power than Student’s t-test. We did not do any simulations to assess power upon correction for multiple testing (Benjamini-Hochberg correction).

### Polysome Profiling

Cycloheximide (CHX; Sigma, G7698) was added at a final concentration of 100μg/ml to muscle cell culture (MB or MT) for 15 minutes. After incubation, plates were placed on ice washed twice with ice-cold DPBS (Lonza, BE17-512Q) containing CHX. Cells were lysed in polysome lysis buffer (20mM Tris-HCl, pH = 7.5 (Sigma, T294), 100mM NaCl (Sigma, 71386), 10mM MgCl_2_ (Sigma, 63069), 1% Triton X100 (Sigma, T8787) was pre-prepared, and 2mM DTT (Sigma, 646563), 100μg/ml CHX (Sigma, G7698), 400U RNAsin plus RNAse Inhibitor (Promega, N261B), 20U Turbo DNase (Ambion, AM2238), cOmplete mini EDTA- free protease inhibitor (Roche, 11836170001). Lysate were homogenised through a 23G needle (Braun, 4657640) and debris removed by centrifugation before optical density (OD) measurement (A_260_) with Nanodrop2000 (Thermo Scientific), aliquoting and snap freezing.

Linear sucrose gradient (10-50%) was prepared following the manufacturer’s instructions (Biocomp Gradient Master). Briefly, 10% sucrose gradients-1X Polysome gradient buffer (50mM Tris-HCl, pH7.5 (Sigma T2194), 50mM NH_4_Cl (Sigma, 09718), 12mM MgCl_2_ (Sigma 63069), 100μg/ml CHX (Sigma, G7698), 0.5mM DTT (Sigma, 646563), 10μL SuperaseIN (Invitrogen, AM2696)- and 50% sucrose gradients-1X Polysome gradient buffer (50mM Tris-HCl, pH7.5 (Sigma T2194), 50mM NH_4_Cl (Sigma, 09718), 12mM MgCl_2_ (Sigma 63069), 100μg/ml CHX (Sigma, G7698), 0.5mM DTT (Sigma, 646563), 10μL SuperaseIN (Invitrogen, AM2696)- were prepared. Open-top thin wall ultra-clear round bottom tube (14x89mm, Beckman Coulter, 331372) were used to load 10% and 50% sucrose gradient with a gradient master 108 (Biocomp). Equal A260 OD of cell lysates were loaded on the sucrose gradients and centrifuged at 4°C at 35000rpm for 3hrs (rotor SW-41/Ti, Beckman Coulter). Polysome profiles were obtained at 254nm wavelength using piston gradient fractionator (Biocomp) and fractions were collected using a Gilson fraction collector and were immediately snap-frozen.

### TCA Precipitation

Fractions corresponding to 1) 40S-60S subunits, 2) 80S ribosomes, 3) light and heavy polysomes were pooled in the same tube and resuspended with 15% (final) of trichloroacetic acid solution (TCA, T0699 Sigma-Aldrich) and precipitated overnight at 4°C. Proteins were further precipitated by centrifugation at 20000xg for 30min at 4°C, washed with 95% Acetone (Merck-Millipore), and centrifuged at 20000xg for 10min at 4°C. Pellets were vacuum-dried at 30°C for 5min using Concentrator Plus (Eppendorf) to remove residual acetone and resuspended in RIPA buffer before Western Blot analysis.

### Imaging

Briefly, cells were grown on coverslips, fixed in 3% paraformaldehyde (EMS, #15714-S)/1% sucrose in 1X PBS pH7,4 for 10min at 37°C before permeabilisation using 0.2% Triton (Trion X100, sigma) for 4 minutes at room temperature. Coverslips were rinsed 3 times with 1X PBS before overnight incubation with primary antibody in 1%BSA (sigma A9647) 1% normal goat serum (ab7481, Abcam) in 1X PBS. After 3 washes the coverslips were incubated with the appropriate secondary antibody (table 3). DNA was stained with Hoechst 33342 (Thermo Fisher Scientific, H1399). Coverslips were mounted with Vectashield medium (Vector Laboratories, H-1000) and images were acquired on a widefield Leica Thunder microscope (processed with LAS X Life Science Software) or a Zeiss LSM 710 scanning confocal microscope (processed with ZEN 2.0 software).

### Gene Ontology (GO) analysis

The ClusterProfiler R package version 3.18.1 was used for all the GO term analyses reported in this study.

## RESULTS

### Generation of a functional eIF3f-BioID1 chimera

Our first objective was to test whether eIF3f interactors can be identified by the biotin ligase-based proximal labelling method. We adapted the method described by Roux et al.^24^, where a protein of interest was fused to *E. coli* biotin ligase mutant protein BirA(R118G) denoted as BirA*. In the presence of biotin, BirA* biotinylates interactor proteins at lysine residues that are within 10 nm distance. Biotinylated proteins thus produced can be immunoprecipitated with the help of streptavidin-coated magnetic beads, and analysed by mass spectrometry (figure 1A). In our study, we fused eIF3f to the N-terminus of BirA*, which was HA-tagged at the C-terminus, using the BirA(R118G)-HA plasmid (referred to as eIF3f-BioID1 in this manuscript). After overexpressing the construct in HEK293 cells, we first assessed the expression of the eIF3f-BioID1 chimera. Detection of an expected, 80kDa band in Western blots by an anti-HA antibody (figure 1B) confirms the expression of the eIF3f-BioID1 chimera. To further assess the functionality of the chimera, eIF3f-BioD1-overexpressing HEK293 cells were grown in presence of 50µM biotin for 24hrs (optimal incubation time determined for BirA*, data not shown). Total lysates were separated by gel electrophoresis and blotted for biotin using streptavidin-coupled horseradish peroxidase (figure 1C). Expression of BirA* alone leads to a strong accumulation of biotinylated proteins in the presence of biotin as shown by the smear, whereas its fusion to eIF3f in the eIF3f-BioD1 changes the pattern of cellular protein biotinylation (figure 1C). Of note, the trace amount of biotin present in the media is not sufficient to induce detectable biotinylation compared to the biotin addition (50 µM, figure 1C). We therefore performed immunoprecipitation and mass-spectrometry as described in the Methods section, to identify the interactors of eIF3f. 184 proteins were more abundant in eIF3f-BioID1-expressing compared to control cells (log2 fold-change (log2FC) > 0, supplementary figure 1A), and 35 out of 184 proteins were significantly enriched (p<0.01, table 4) among which eIF3f exhibited the strongest enrichment (log2FC=7.27), and the set included 6 other subunits of the eIF3 complex (figure 1D). Importantly, all the proteins identified from cells expressing eIF3f-BioID1 were absent in pull downs from cells expressing only the BirA* protein (supplementary figures 1C and D). Gene set enrichment analysis showed a highly connected network of enriched proteins (figure 1E) that included members of the eIF3 complex (subunits a, d, c, g, h, l and f). Interestingly, proteins involved in the regulation of mRNA stability were also enriched (supplementary figure 1B). Altogether these results demonstrate that the eIF3f-BioID1 chimera is functional in the assembly of the eIF3 complex.

**Figure 1:**
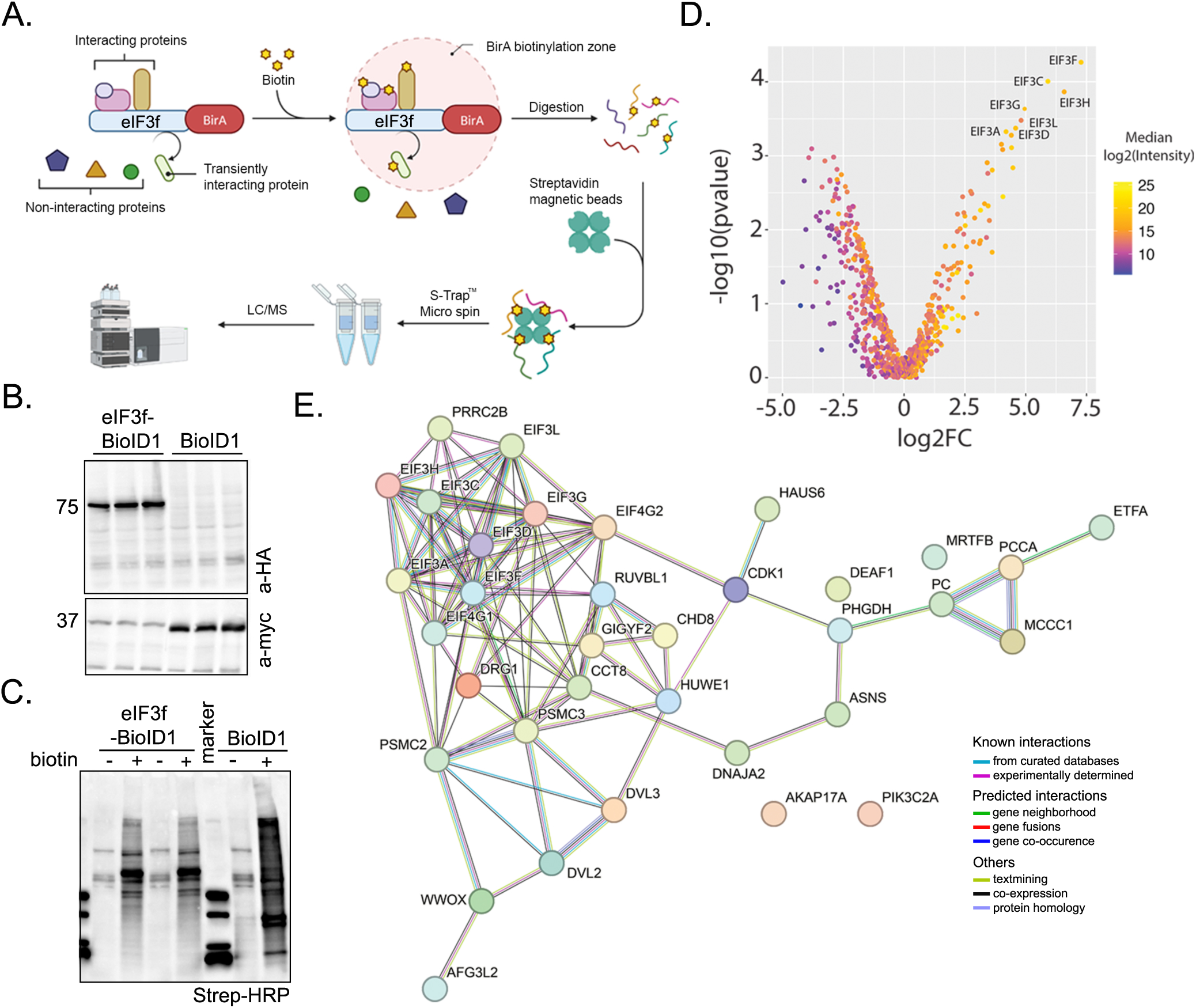
Expression and characterization of a functional eIF3f-BioID1 chimera. **A.** Workflow for the LC/MS identification of eIF3f-BirA (eIF3f-BioID1) proximal interactors *in cellulo* with proximity-dependent biotin labelling. Created in BioRender. **B.** Immunoblot detection of eIF3f-BioID1 chimera expression. Lysates from transfections of HEK293 cells (triplicates) were separated on SDS gel and expressed proteins were detected with indicated antibodies. **C.** 24hrs upon transfection (as described in B), cells were cultured in presence or not of 50µM biotin for additional 24hrs before lysis. Immunoblot blot detection of biotinylated proteins (streptavidin-HRP) upon expression of eIF3f-BioID1 or BioID1 shows their respective specificity. **D.** Volcano plot distribution of the proteins identified by LC/MS upon streptavidin-dynabeads purification of biotinylated proteins from transfected HEK cells. **E.** Direct (physical) and indirect (functional) interaction map of eIF3f interacting proteins identified with high confidence.

**Table 4.**
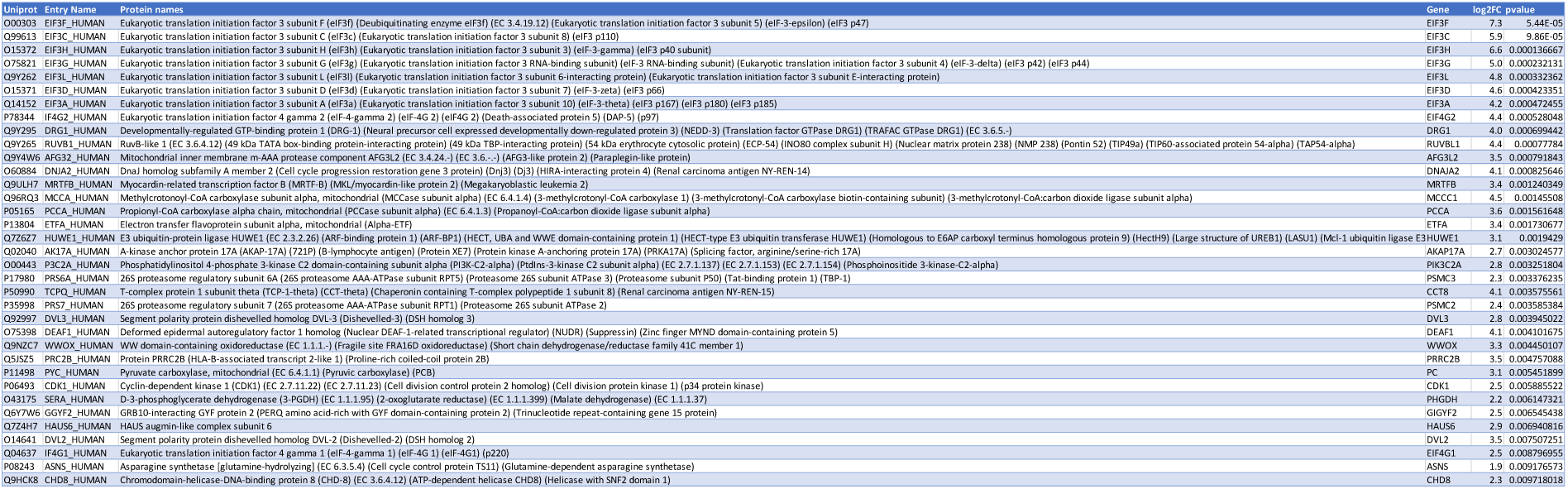
eIF3f-BioID1 interactors in HEK293 cells.

### eIF3f-BioID1 chimera is more efficient than eIF3f-BioID2 *in cellulo*

Before creating a stable cell line expressing eIF3f-BioID1 from the eIF3f locus, we explored the efficacy and functionality of other fusion constructs. As the BirA* ligase (BioID1) is relatively larget, 35kDa, we tested a smaller, recently described Biotin ligase, BioID2 (27kDa). BioID2 lacks of the DNA-binding domain and has been shown to display more efficient and specific biotin transfer activity^31^. We also assessed the influence of N-terminal (NH2 or N-ter) versus C-terminal (COOH or C-ter) fusion of the biotin ligase to the eIF3f protein. Both (N-ter and C-ter) BioID2 chimeras were transiently expressed in HEK293 cells. Western blot analysis revealed a 70KDa band migrating below the previously described band representing eIF3f- BioID1 (supplementary figure 2A). Upon expression of N-ter and C-ter tagged BioID2 chimeras and biotin provision, biotinylated proteins were detected by Western blot with streptavidin-HRP conjugate. As shown in supplementary figure 2B, the BioID2 enzyme alone displays a strong biotinylation activity. The BioID2 fused to the eIF3f N-terminus appeared mislocalised (accumulation in giant cytoplasmic vesicles, data not shown), and gave a strong biotinylation pattern accompanied by a strong band at low molecular weight (∼30KDa) (supplementary figure 2B), which could result from a cleavage of the self-biotinylated fusion protein. Therefore, we selected the eIF3f C-ter fused BioID2 chimera (eIF3f-BioID2) to further proceed with MS/MS coupled streptavidin pull down (supplementary figures 2 C and D). Among the 118 proteins enriched after the streptavidin pull down (log2FC>0), only 18 had a p-value <0.01 (supplementary figures 2 E; table 5) and 11 of these were already identified in the eIF3f_BioID1 pull down assay. Thus, the sensitivity of partner identification was higher with BioID1, which also appear to be devoid of (cleavage products of the fusion protein).

**Table 5.**
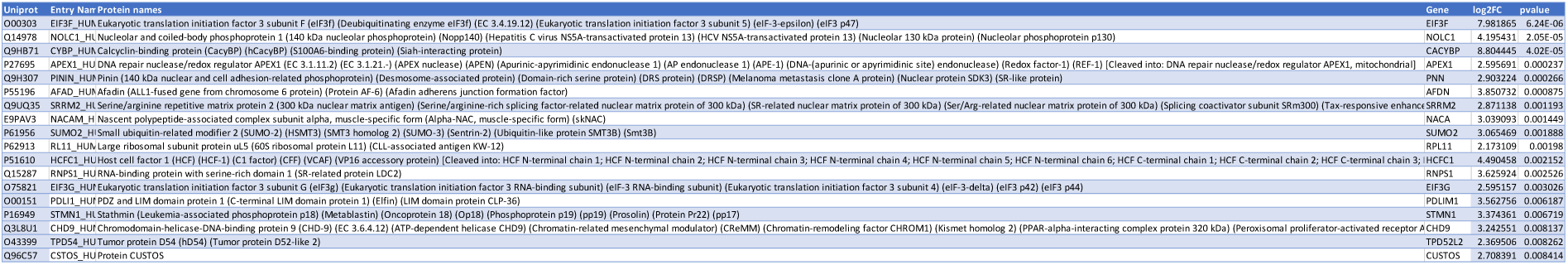
eIF3f-BioID2 interactors in HEK293 cells.

### Generation of genetically engineered EIF3F-BioID1 knock-in a human muscle cell line

Based on the above-described analyses of various BioID tags, we decided to use BioID1 to endogenously tag eIF3f at its C-terminus in immortalised human muscle myoblasts. We used the CRISPR-Cas9 system to genetically engineer the EIF3F locus. Figure 3A shows sgRNAs and homologous arm designing strategy for integration of BioID1 at the eIF3f C-terminus. To construct the cell line, we used two plasmids, expressing Cas9 and either sgRNAs sg1 or sg2 to target the EIF3F exon 10 flanking regions (figure 2A). The pSpCas9(BB) plasmid backbone was used to subclone sgRNAs with each sgRNA assigned to either GFP- or mCherry-expressing plasmid to allow sorting of double positive (DP) cells expressing both sgRNAs (figure 2B). To integrate BioID1 in the EIF3F coding frame by homologous recombination, the donor plasmid contained: left homology arm (HLA; complementary to exons 8 and 9), exon 10, HA-tagged BioID1, T2A “cleavable” peptide and blue fluorescent protein (BFP; used for fluorescent activated cell sorting of DP cells) with a stop codon, and, finally, a right homology arm (HRA; complementary to the 3’ UTR of *EIF3F* gene) (figures 2A and supplementary figure 3A and B). Following the selection for cells expressing both sgRNAs, the DP cells were grown for 48 hrs before a second FACS selection of BFP positive (for construct integration) myoblasts (figure 2B). After expansion, the single cell clones (sc) were screened for donor DNA insertion by PCR amplification of the 3’ region of the EIF3F locus (figure 2C) and selected positive clones were further confirmed by sequencing of the PCR product (see supplementary figure 3 for the sequence). As shown in figure 2B, all selected sc clones were heterozygous for EIF3F- BioID1 insertion. However, single allele integration was sufficient to detect the chimeric protein expression in the selected single cell clones 4 and 6 (later referred as sc4 and sc6), in which anti-eIF3f, anti HA and anti-BirA antibodies all showed reactivity towards a ∼80kDa protein. This was absent from the parental cell lysate. Furthermore, the expression of the endogenous eIF3f was not significantly altered (figure 2D). In conclusion, we successfully constructed knock-in cell lines expressing the eIF3f-BioID1 chimera (figure 2A; supplementary figure 3C).

**Figure 2:**
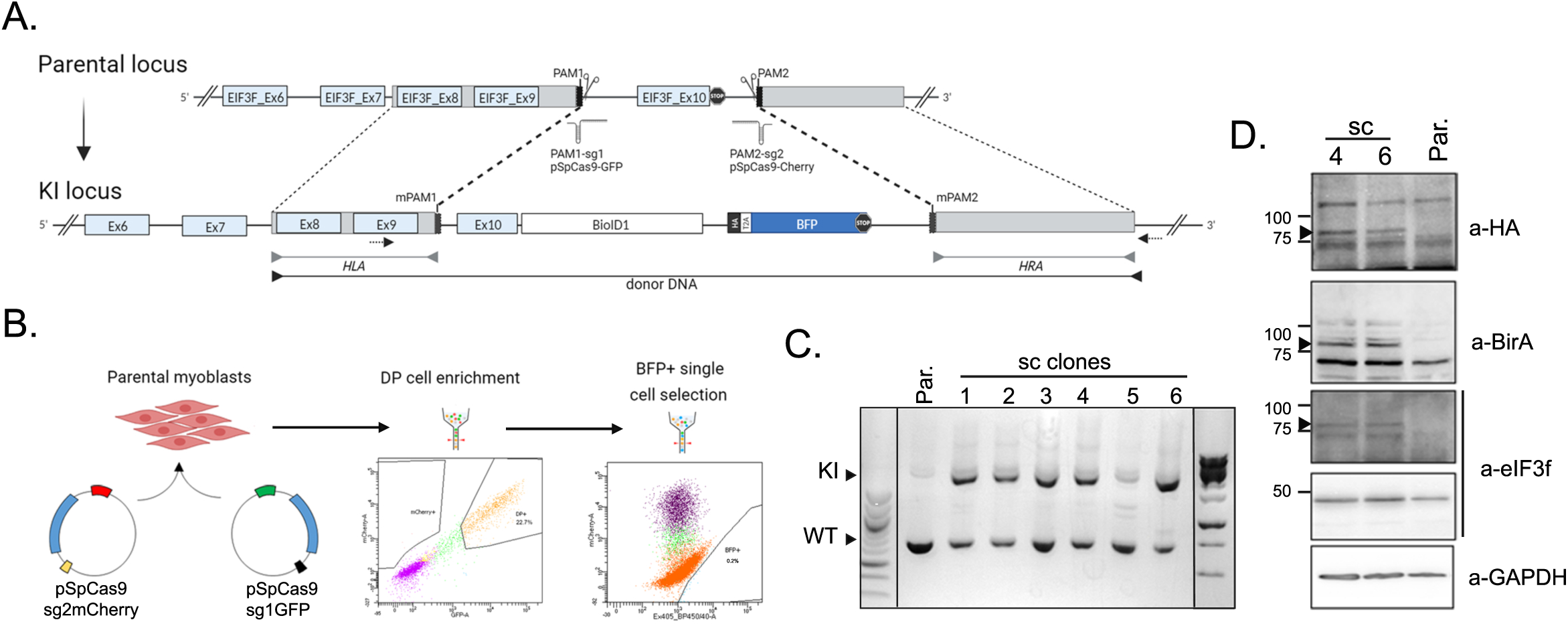
Generation of EIF3F-BioID1 knock-in human muscle cells. **A.** Schematic representation of the Cas9 mediated establishment of EIF3F-BirA knock-in cells. Two small guide RNAs were used to remove EIF3F Ex10 and stop codon allowing the insertion of the donor DNA sequence consisting of the EIF3F Ex10 fused to the BirA and HA sequences together with a cleavable (T2A, self-cleaving peptide) blue fluorescent protein (BFP) flanked by left and right homology arms. Arrows show localization of forward (EIF3F_scrn_F) and reverse (EIF3F_scrn_R) primers (see table 2) used for PCR validation of the single cell clones isolated by FACS sorting (BFP^+^). Created in BioRender. **B.** Graphical illustration of EIF3F-BioID1 myoblast single clone selection. pSpCas9 plasmids expressing a pair of sgRNAs (sg1 and sg2) where each sgRNA was tagged by either GFP (green box) or mCherry (red box) fluorescent protein were electroporated into myoblasts. Upon double positive (DP) population selection, single BFP positive cells were sorted into 96 well plates. **C.** DNA extracted from each single-cell colony was purified and subjected to PCR with specific primers distinguishing between wild type (WT) and knock-in (KI) alleles at the EIF3F locus. **D.** Cell lysates extracted from proliferating single cell clones (sc4 and sc6) and parental cells were subjected to Western blotting and chimer protein expression eIF3f-BioID1 (80KDa) was detected using antibodies directed against HA and/or BirA.

**Figure 3:**
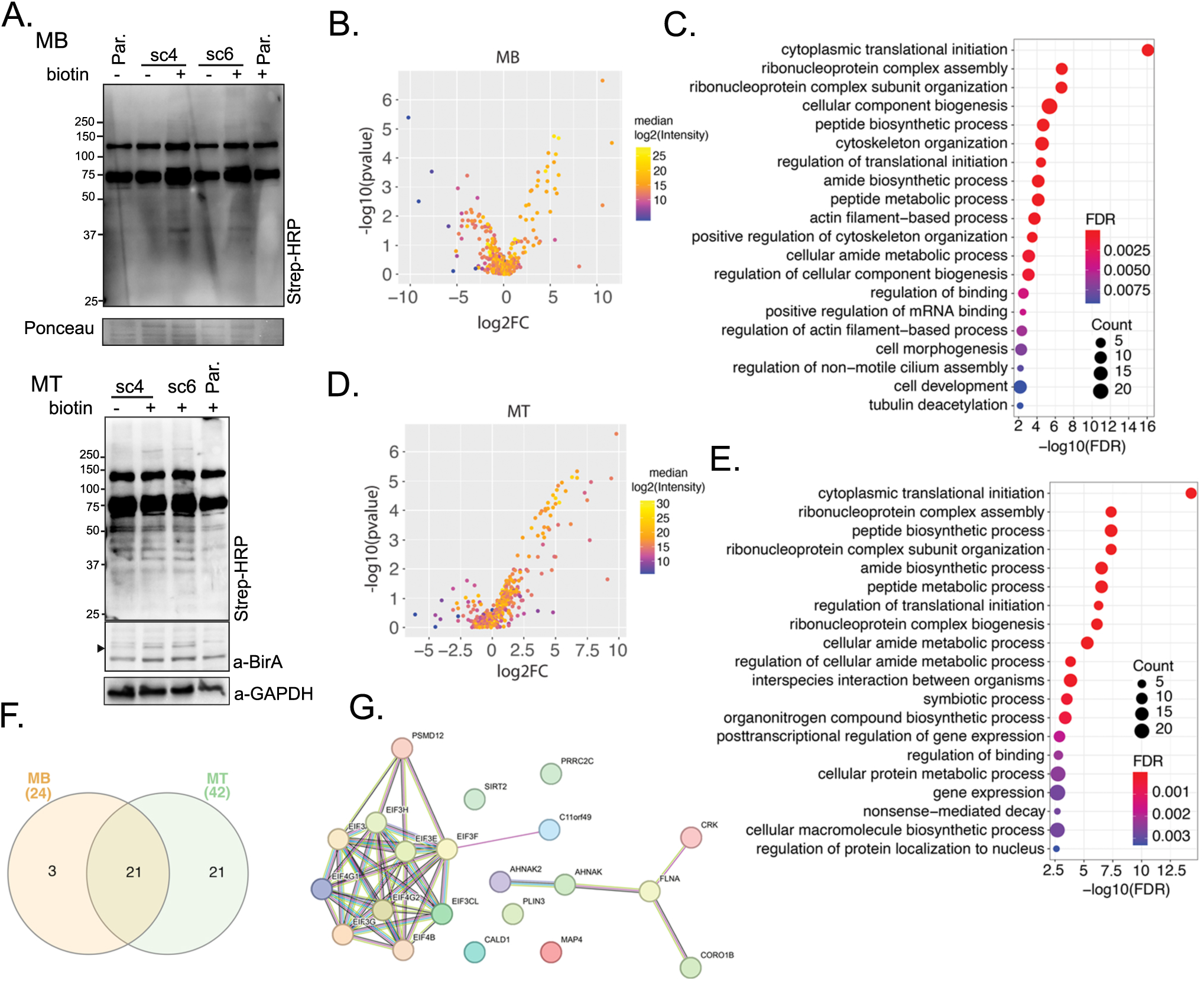
eIF3f interacts with the core translational machineries in both MB and MT. **A.** Expression and functionality of eIF3f-BioID1. Single cell clones 4 and 6 of proliferating MB and differentiated MT were cultured in the presence or absence of 50μM biotin for 24hrs. Endogenously biotinylated proteins were visualised with streptavidin coupled HRP in both MB and MT. Expression of eIF3f-BioID1 chimer is shown in MT (anti-BirA). GAPDH or Ponceau staining were shown as loading control. **B.** Volcano plot distribution of streptavidin-dynabeads immobilised proteins (biotinylated proteins) identified by LC/MS, from proliferating (myoblasts, MB) sc4/6 cultured in presence of biotin versus parental cells (supplementary figure 4A and B). **C.** Functional enrichment analysis of proteins identified in B (p<0.01 and log2FC>0). Only the top 20 pathways (based on FDR) are shown. The same analysis shown in **D** and **E** was carried out with differentiated (day 6, myotubes, MT) sc4/6 cultured in presence of biotin versus parental cells. **F.** Venn diagram representation of identified eIF3f binding partners from MB and MT. **G.** Interaction map of the 21 identified binding partners of eIF3f-BioID1 from the Venn diagram intersection between MB and MT.

### Identification of a shared proximal interactome of eIF3f in MB and MT

To interrogate the role of EIF3F in skeletal muscle cell physiology, we employed the generated EIF3F-BioID1 single muscle cell clones to identify the endogenous eIF3f interactors in proliferating myoblasts (MB) and in post-mitotic myotubes (MT). To this end EIF3F-BioID1 clones (sc4 and sc6) and non-engineered parental (Par.) cells were cultured in the presence of biotin (supplementary figures 4A and B). Obtained samples were lysed, the lysates were subjected to gel electrophoresis and blotted for biotinylated proteins using streptavidin-HRP (figure 3A). Figure 4A shows protein biotinylation in cells cultured in the presence of biotin. Aliquots of respective sample lysates were used for the pull-down purification of biotinylated proteins with streptavidin coupled Dynabeads. MS-MS analysis identified 65 proteins that were more abundant (log2FC>0) in MB cells expressing eIF3f-BioID1 when compared to Par. cells. Among these 65 proteins 24 were significantly enriched (p<0.01) (figures 3B and C; table 6; supplementary figure 5A). In MT, 42 proteins out of 260 were significantly enriched (log2FC>0, p<0.01) (figures 3D and E; table 7; supplementary figure 5B). Among the 24 binding partners of eIF3f in MB, 21 were also identified as binding partners in differentiated MT (figure 3F; table 8). The vast majority of these 21 interactors (figure 3G) are involved in translation initiation, regulation of translation initiation complex assembly as well as regulation of RNA binding (figures 3C and E; supplementary figure 5C). They include eIF3 complex subunits a, cl (c-like), e, g, and h, as well as eIF4B and eIF4G1 and eIF4G2 (figure 3G). The identification of well-established interactors of eIF3f support the functionality of the chimera in both MB and MT. A second cluster of interacting proteins (figure 3G) is related to the cytoskeleton myofibril and actin filament/cilia organisation (figure 3C), e.g. Microtubule-binding proteins 4 and 1A (MAP4; MAP1A), Microtubule actin crosslinking factor 1 (MACF1), Septin9, ALPK3, and Centriolar satellite-associated tubulin polyglutamylase complex regulator 1 (CSTPP1). Together, these observations suggest that the main function of eIF3f in skeletal muscle cells is the regulation of protein synthesis and that eIF3f activity could be spatially restricted to specific subcellular compartments such as the Z-disc and the centriole. Only 3 proteins, the myocardin-related transcription factor B (MRTFB), the LIM domain-only protein 7 (LMO7) and the eukaryotic translation initiation factor 5 (eIF5) were specifically found in MB. In contrast, along with 19 other proteins, eIF3 complex subunits l and m were identified only in MT. The observed differences between eIF3f interactors in MB and MT cells may indicate the role of eIF3f in translational regulation changes during the differentiation process.

**Figure 4:**
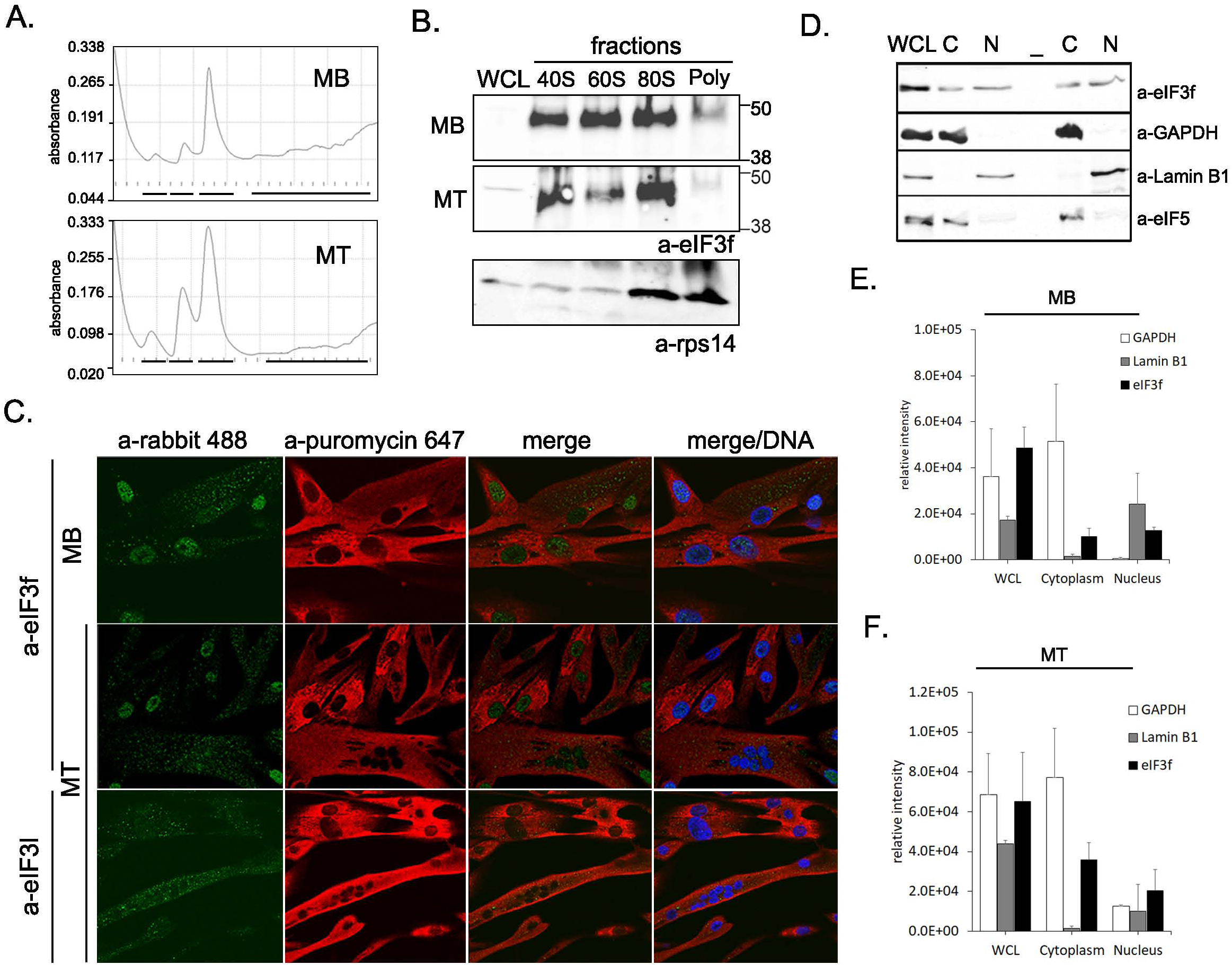
eIF3f subcellular distribution and related functions. **A.** Polysome profiles generated from MB vs MT parental cells. The same amount of lysate (OD 260) was loaded on a sucrose gradient (10-50%) before collection and pooling of 40S, 60S, 80S and polysome fractions as represented by black lines respectively from left to right on the bottom of each profile. **B.** The proteins from the 4 collected fractions were precipitated before separation and immunoblotted with the indicated antibodies (WCL, whole cell extract). Ribosomal protein rps14 is used as a marker. **C.** Subcellular distribution of “active sites of protein synthesis” in human muscle cells (parental) MB and MT as visualised upon pulse chase incorporation of puromycin (SunSET method^25^) by confocal microscopy with anti-puromycin coupled AF647 (red). Both eIF3f and eIF3l proteins were detected with respective antibodies and visualised with Cy3 coupled anti-rabbit secondary antibodies. DNA was stained with Hoechst. **D.** Whole cell lysate (WCL) or fractionated (C: cytoplasm and N: nuclear) lysates from parental MT were resolved on SDS/PAGE and probed with anti eIF3f or anti eIF5 antibodies. GAPDH and Lamin B1 were used respectively as cytoplasmic or nuclear markers. Infrared signals (700, 800nm) from the secondary antibody were acquired with Odyssey (Li-Cor). **E.** Relative intensity signals for eIF3f, Lamin B1 and GAPDH from D (parental MB (n=3) are presented and compared with parental MT (n=3). Original immunoblots used for quantification are displayed in supplementary figures 6C and D.

**Table 6.**
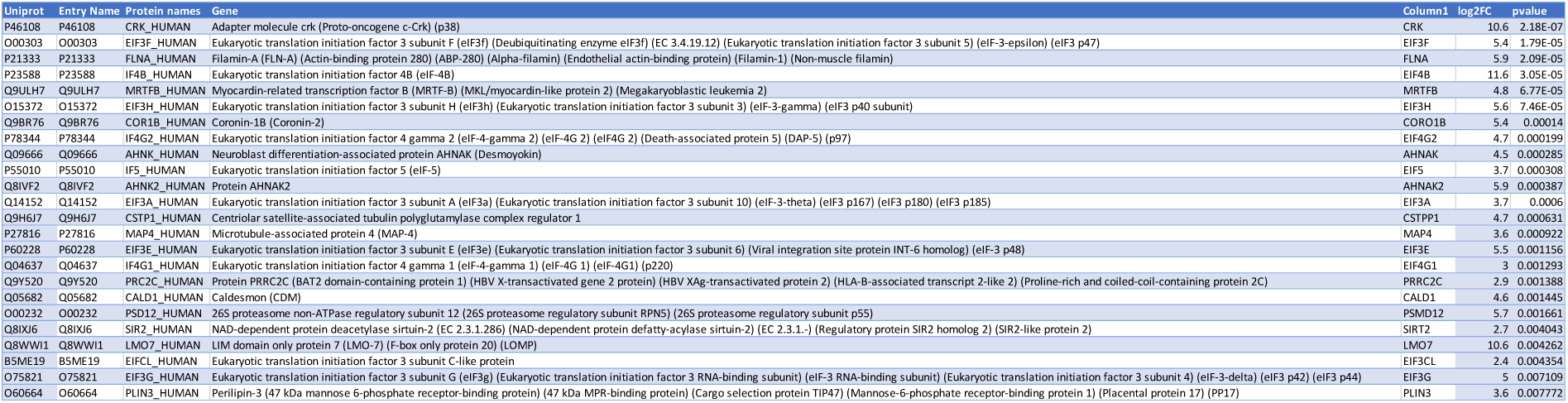
eIF3f-BioID1 interactors in sc4/6 MB.

**Table 7.**
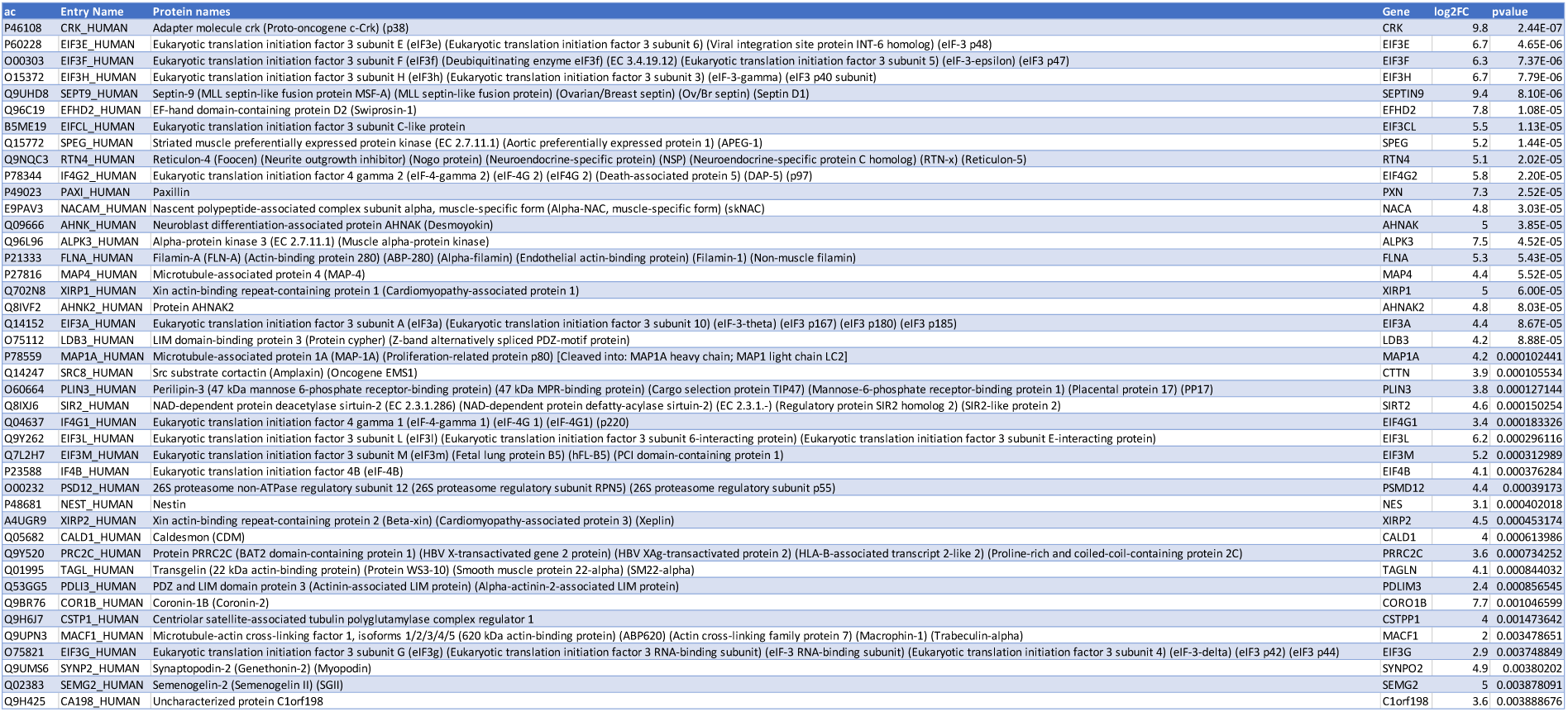
eIF3f-BioID1 interactors in sc4/6 MT.

**Table 8.**
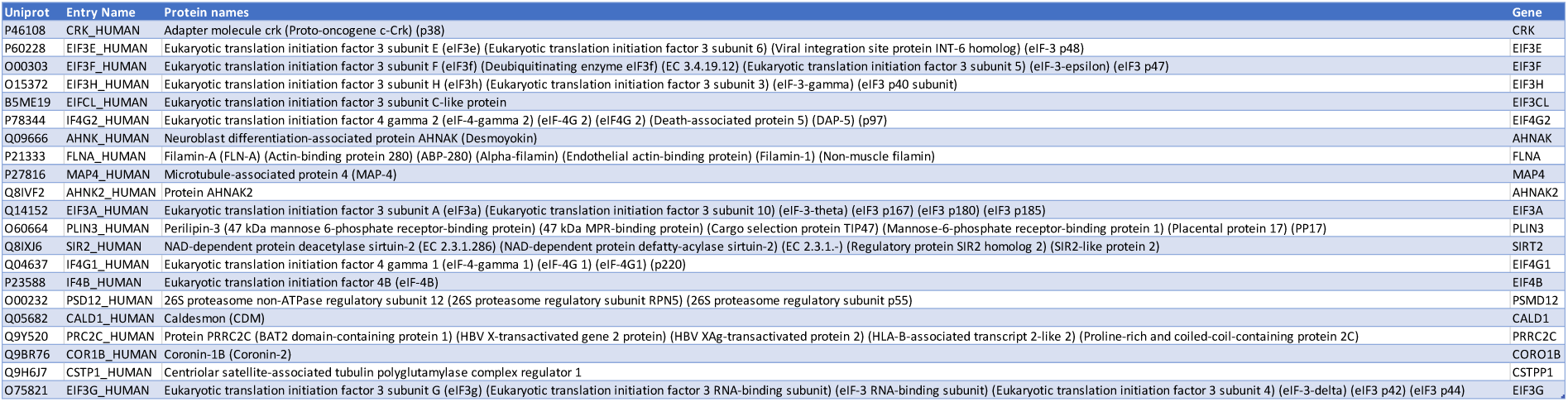
The 21 eIF3f interactors conserved in both MB and MT.

To validate the identified translation-related interactors of eIF3f protein in human skeletal muscle cells, we performed profiling of polysomes isolated from MB and differentiated MT. As shown in figure 4A, polysomes are relatively less abundant in MT in comparison to MB whereas 40S, 60S and 80S peaks are stronger in MT, suggesting reduced level of translation in MT. Further, Western blot analysis showed the presence of eIF3f in 40S, 60S and 80S fractions, supporting its role in translation initiation complex assembly in skeletal muscle cells (figure 4B).

Additionally, as we identified eIF3f binding partners involved in the muscle fibre (Z-disc) and cytoskeleton organisation, we wondered whether the subcellular pattern of eIF3f localisation is related to sites of active translation in skeletal muscle cells. We employed the Sunset method^32^ which involves pulse chase (30min) incorporation of puromycin, an amino-acid analogue which incorporates into newly synthetised proteins at very low concentrations (10μg/mL) and allows the detection of active translation sites in cells by the use of Alexa-Fluor488 coupled anti-puromycin antibodies. Immunofluorescence staining was performed in both cultured human parental MB and MT and confocal microscopy was used to analyse cytoplasmic distribution of puromycin labelled proteins (figure 4C). Whereas the staining in MT displayed a diffuse cytosolic pattern, staining in MB displayed punctuated distribution in regions surrounding the MB nuclei, suggesting association of translation process with the ER/endosomal compartment. Furthermore, we observed higher puromycin staining intensity in MB compared to MT, likely reflecting an increased protein synthesis in MB cells, which is consistent with increased abundance of polysomes in gradient analysis profiles of MB cells (figure 4A).

To validate that eIF3f is present in these “nascent” translation sites, we co-stained MB and MT cells with an anti-eIF3f antibody. The cytoplasmic eIF3f co-localised with the puromycin labelled proteins (figure 4C), as did eIF3l, another subunit of the eIF3 complex (figure 4C). Taken together, these observations support the role of eIF3f in active translation in both MB and MT.

Surprisingly, the co-staining with anti-eIF3f antibody also showed eIF3f localisation in the nucleus (figure 4C) in both MB and MT. This indicates a nucleus-specific function of eIF3f. We also verified nuclear staining of eIF3f both in MB and MT without puromycin co-staining (supplementary figures 6A and B). To confirm the result of immunostaining, we carried out eIF3f immunoblotting of the whole cell lysate, as well as nuclear and cytoplasmic fractions of cells. The immunoblot detected endogenous eIF3f protein in whole cell lysates, nuclear (N) and cytoplasmic (C) fractions from both MB and MT (figure 4D and supplementary figures 6C and D). The purity of fractions was confirmed by blotting for the nuclear marker Lamin B1 and the cytoplasmic marker GAPDH (figure 4D). The result showed that the nuclear fraction was not contaminated with the cytoplasmic fraction and vice versa. Western blot quantification of eIF3f protein showed the presence of the protein in both cytoplasmic and nuclear fractions of muscle cells (figure 4E and F).

Altogether, these observations demonstrate a dual (cytoplasmic and nuclear) localization of eIF3f in both MB and MT. In the cytoplasm, eIF3f participates in translation in association with other eIF3 subunits, whereas the nuclear eIF3f may be involved in some aspects of gene regulation, as nuclear proteins (according to https://www.genecards.org/nomenclature), such as the transcription factor MRTFB and regulators Sirt2, CHD8, and RUVLB1, were found among the eIF3f interactors (supplementary figure 6E).

### eIF3f interacts with the vesicle marker LAMP1

We found that eIF3f strongly interacted with the components of eIF3 complex in muscle cells and that 4 of them, namely eIF3a, eIF3cl, eIF3e and eIF3h belong to the 21 common proteins found both in MB and MT (figure 3F). Together with eIF3f, these subunits belong to the conserved octamer PCI/MPN structural scaffold which functions in different steps of translation^6^. To identify interactors that could differentially regulate the function of these five “core eIF3f-eIF3” proteins (figure 5A) in terminally differentiated cells, where eIF3f was reported to promote hypertrophia, we normalised all protein intensities using intensities of the five “core eIF3f-eIF3” proteins as a reference. We then analysed the differential association of the identified interactor proteins (figures 3B and D) in MT relative to MB. We identified 25 proteins, 18 of which were significantly enriched for their interaction with the “core eIF3f- eIF3” in MT (supplementary figure 7A), and some being known to localize to specific subcellular compartments. These are the lysosome-associated protein LAMP1, the ER associated protein reticulon 4 (RTN4), the sarcomere associated proteins XIRP1/2, LDB3 and the MYH3/8. Since LAMP1 was the top-enriched interactor of the “core eIF3f-eIF3” in MT, we interrogated the molecular interaction between endogenous eIF3f and LAMP1 proteins in the parental MT lysate by immunoprecipitation. We found that LAMP1 was co-immunoprecipitated with anti-eIF3f antibody and, reciprocally, eIF3f was pulled-down by the anti-LAMP1 antibody (figure 5B). This result strongly supports the interaction of eIF3f with LAMP1 in MT. In immunofluorescence experiments, we observed a punctae/aggregate-like distribution of eIF3f in the cytoplasm (figure 4C) and employed immunofluorescence to visualise the localization of LAMP1 in MT cytoplasm. LAMP1 was distributed throughout the cytoplasm with a pronounced perinuclear staining and a characteristic endosome-like pattern (supplementary figure 7B). Importantly, both eIF3f and LAMP1 antibodies stained small aligned aggregate-like structures in the cytoplasm of MT (figure 5C). Altogether these data suggest that eIF3f may be associated with specific “translation spots” at LAMP1-positive late endosomes of MT cells.

**Figure 5:**
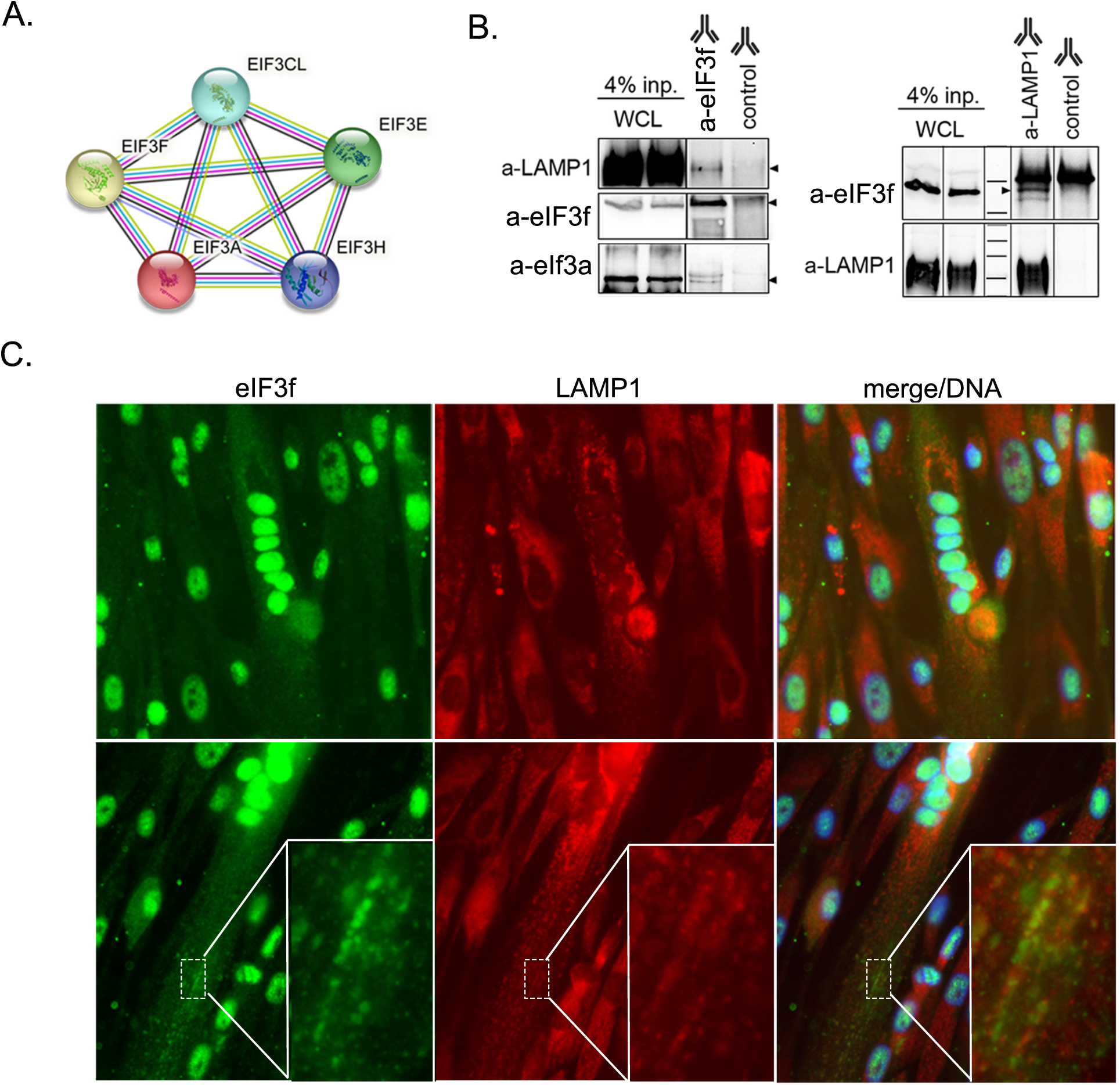
Physical interaction between eIF3f and the autophagy vesicle marker LAMP1. **A.** Protein-protein functional enrichment of the “core eIF3f-eIF3” used for normalization and identification of positively enriched interactors in MT **B.** Western blot detection of eIF3f and LAMP1 after immunoprecipitation from parental MT (day 6) lysates. Anti-eIF3f or anti-LAMP1 antibodies were used for precipitation as indicated and corresponding isotype was used as control. eIF3a protein detection was used as a positive control for anti-eIF3f immunoprecipitation and 4% input whole cell lysate (WCL) was used as an internal control for protein detection. **C.** Immuno-detection of eIF3f and LAMP1 visualised with anti-rabbit Cy3 and anti-mouse Cy5 coupled secondary antibodies respectively in parental differentiated (day 6) myotubes (MT). DNA was stained with Hoechst. Dashed rectangles are displayed in a magnified rectangle to appreciate co-localisation of both proteins in the cytoplasmic part of MT.

## DISCUSSION

eIF3f was recently proposed to be an essential factor for skeletal muscle growth and maintenance, though the underlying molecular mechanisms are not known. To enable a better understanding of the role of eIF3f in muscle cells we here used the proximity labelling approach to provide the first analysis of the eIF3f protein interactome. We generated an eIF3f-BioID1 chimera whose subcellular distribution is similar to that of the endogenous protein and that additionally displays biotin ligase activity. Analysis of mass spectrometry data obtained from HEK293 cells transiently expressing the eIF3f C-ter tagged chimera (eIF3f-BioID1) revealed known partners of eIF3f, indicating that the presence of the BioID1 tag at the C-terminus did not interfere with eIF3f interaction with its partners (figure 1).

We then created a stable human muscle cell line by genomically tagging EIF3F with BioID1 using CRISPR/Cas9 (figure 2). The cell line was heterozygous in terms of BioID tagging but the tagged eIF3f was sufficient to identify eIF3f interactors, even in the presence of endogenous eIF3f. The interactome derived from proximity labelling was enriched in known eIF3f interactors in both MB and MT cells, as expected. 21 proteins were found to interact with eIF3f in both muscle myoblasts and myotubes (see table 8), and most of these were related to translation initiation complex assembly (the core subunits a, cl, e, g, h of the eIF3 complex, the initiation factors eIF4B, eIF4G1 and G2, figure 3). Further, eIF3f was also found to interact with the PRRC2 protein, which is involved in promoting translation of mRNAs containing upstream open reading frames (uORFs) via leaky scanning^33^. PRRC2 has three isoforms; PRRC2A, B and C, all representing core components of stress granules^34^ and we found PRRC2B in both MB and MT.

The eIF3f-BioID1 chimera also identified cell-stage-specific interactors of eIF3f (e.g. l and m subunits of eIF3f in MT), suggesting differential regulatory role of eIF3f in MB and MT. We identified markers for specific subcellular compartments such as the ER associated protein reticulon 4 (RTN4) and the sarcomere associated proteins XIRP1/2, LDB3 and the MYH3/8. We found several components of the cytoplasmic structures as proximal interactors of eIF3f, suggesting a possible role of eIF3f in localised translation. For example, the PDZ and LIM domain 3 (PDLIM3) is a cytoskeletal protein that co-localizes with α-actinin on the Z line and ALPK3 is a myogenic kinase located at the M band of the sarcomere of the striated muscle where it interacts and phosphorylates MAP4^35^. We also identified a muscle-specific isoform of the nascent polypeptide-associated complex subunit alpha (skeletal NACA (skNAC)), which is required for myofibril organization^36^. Remarkably, regulators of microtubule dynamics and cleavage (ALPK3 and CSTPP1), microtubule-associated proteins (MAP4, MAP1A, MACF1), the actin binding and crosslinker protein (EFDH2/swiprosin) and Septin9 also emerged as eIF3f interactors. CSTPP1, already predicted *in silico* to bind eIF3f, is a new member of the tubulin glutamylase family that was recently shown to orchestrate the microtubule and actin assembly in the regulation of nuclear shape^37,38^. Tubulin glutamination regulates MAP1A binding and vesicle transport in synapses, while the spectraplakin scaffold protein MACF1 acoordinates the crosstalk between actin microfilaments and microtubules. MACF1 was recently shown to control the microtubule-dependent localization of extra-synaptic myonuclei and mitochondrial biogenesis in mouse muscle^39^. Moreover, several of these proteins display a centriolar satellite localisation (CTPP1, MAP4, MAP1A, Septin9) and were already shown to interact with RNA^40^. These observations led us to hypothesize that the cytoplasmic eIF3f function in translation might be locally restricted to specific sites/spots defined by these proteins. To validate the specific localization of eIF3f, we visualized translation in both MB and MT with the sunset method (figure 4). Puromycin staining was distributed homogeneously in the cytoplasm in MT, while showing cytoplasmic punctae/aggregates in MB. These punctae co-stained with anti-eIF3f antibodies.

To further identify cell stage specific binding partners that could modulate the functionality of eIF3f in translation, we defined a “core eIF3f-eIF3” composed of eIF3a, eIF3cl, eIF3e, eIF3h and eIF3f, and determined the intensity of interactor signal relative to the intensity of the five “core eIF3f-eIF3” proteins. LAMP1 was the top enriched interactor with the “core eIF3f-eIF3” in human myotubes. LAMP1 is a type 1 transmembrane protein identified as the major constituent of the lysosomal membrane involved in autophagosome formation (for review^41^) and late endosome trafficking^42^. Aside from its function in the lysosomal degradation pathway, LAMP1 together with Rab7 were identified as late endosome markers associated with active local translation and responsible for the delivery of the newly synthesised proteins to mitochondria in neurons^43^. We could validate the interaction of endogenous eIF3f and LAMP1 proteins by reciprocal immune-affinity pull-downs (figure 5). Immunofluorescence analysis of the LAMP1 subcellular distribution revealed a “granular” pattern in the cytoplasm with some granules/puncta being also positive for eIF3f staining.

Surprisingly, we also identified eIF3f-interacting factors that have a nuclear localization: the transcription factor MRTFB and the DNA binding proteins CHD8, RUVLB1. Consistently, we have observed the presence of eIF3f in nuclei by immunofluorescence and biochemical fractionation experiments, in both MB and MT. Such observations were already reported in cancer cells where exogenously overexpressed eIF3f was also in the nuclear fraction^21,44^. The phosphorylation of eIF3f by CDK11 was proposed to drive eIF3f nuclear translocation during induction of apoptosis. Notably, some other essential components of the translation initiation machinery were previously reported to be present in the nucleus (for review ^45^). A considerable fraction of the cap-binding protein eIF4E localizes to the nucleus, where the protein is possibly involved in the export of some cellular mRNAs^46^ (see also supplementary figures 6C and D). A shuttling protein, eIF4E-T, has been identified as a transporter of eIF4E to the nucleus^47^. Finally, ribosome biogenesis factors such as the nucleolar proteins RRS1 (Ribosome biogenesis regulatory protein 1) and TTF1 (Transcription Termination Factor 1) also appeared among eIF3f interactors, which makes it tempting to speculate that eIF3f also plays a role in the ribosome biogenesis process in the nucleus.

In summary, we generated and characterised a muscle cell line expressing a functional BioID - tagged eIF3f and showed that a main function of eIF3f in muscle cells is in the regulation of translation. An interesting question to address is whether eIF3f regulates translation of specific mRNAs and at specific locations to stimulate muscle hypertrophy. Since the E3 ubiquitin ligase MAFbx/atrogin-1 targets eIF3f for degradation during atrophy^2^, one could speculate that preventing eIF3f degradation may rescue atrophy. However, the exact role of eIF3f depletion in atrophy is not known. Our data also raise a possibility that eIF3f is involved in ribosome biogenesis, although other moonlighting functions of eIF3f in the nucleus are also possible.

## Acknowledgements

We thank the Proteomics Core Facility (PCF) of the Biozentrum from the University Hospital of Basel and the Flow Cytometry Facility of the Department of Biomedicine from the University Hospital of Basel for their support and expertise. This work was supported by a grant from the Swiss Foundation for Research on Muscle Diseases (FSRMM).

## Author contributions

L.T. and N.M. conceived and designed the experiments, M.S. and M.Z. supervised the work, L.T., N.M., S.A., and Y.I.E., conducted the BioID experiments and established single cell clones, T.B. performed MS/LC analysis, M.A. performed data analysis, B.E performed confocal microscopy analysis. L.T., N.M., M.Z. and M.S. wrote the manuscript and all authors reviewed the manuscript.

## Data availability statement

All mass spectrometry proteomics data associated with this manuscript have been deposited to the ProteomicsXchange consortium via MassIVE (https://massive.ucsd.edu) with the accession number (Dataset is currently private).

## Competing interests

The authors declare that they have no competing interests.

## Notes

### Competing Interest Statement

The authors have declared no competing interest.

