## Supplemental Figures and Legends for "Identification of new interactors of eIF3f by endogenous proximity-dependent biotin labelling in human muscle cells": Supplementary Fig_Sc reports.pdf

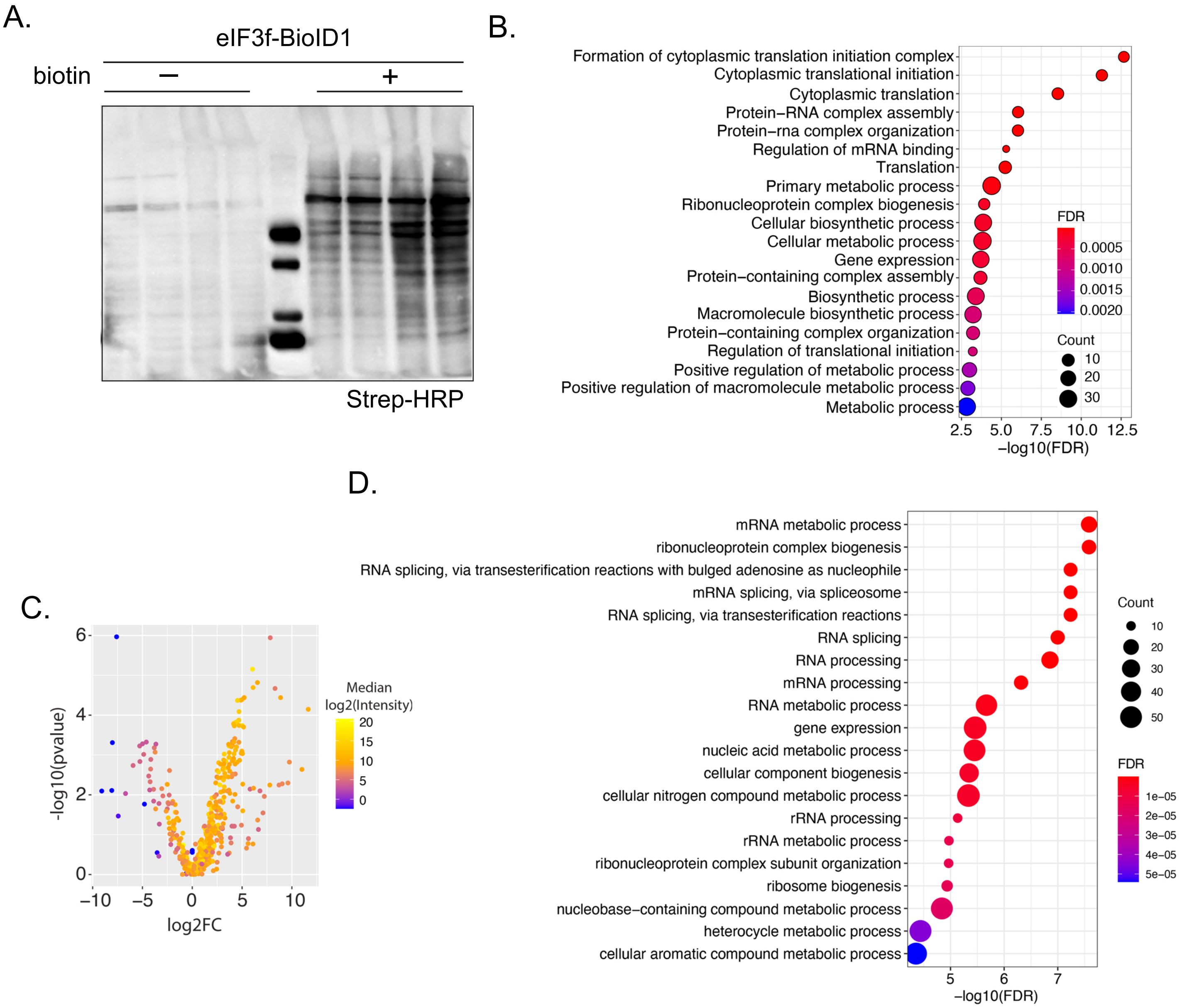


**Supplementary Figure 1: Transient BioID1 biotinylation in HEK293 cells.**

A. Streptavidin coupled HRP (abcam ≠ab7403) detection of biotin incorporation in lysates from HEK293 of overexpressing eIF3f-BioID1 and cultured 24hrs in presence or not (+/-) of 50µM biotin. The same lysates were further used for MS/LC coupled Streptavidin-dynabeads pull down experiment. B. Functional annotation of proteins showing a significant enrichment after pulldown from HEK293 cell lysate expressing or not eIF3f_BioID1 in presence of biotin. Only the top 20 pathways (based on FDR) are shown. C. Volcano plot distribution of the protein identified by MS/LC upon streptavidin-dynabeads purification of biotinylated protein upon overexpression of BioID1 only protein in HEK293 cells. D. Functional annotation of proteins showing a significant enrichment with BioID1 only protein. Only the top 20 pathways (based on FDR) are shown.

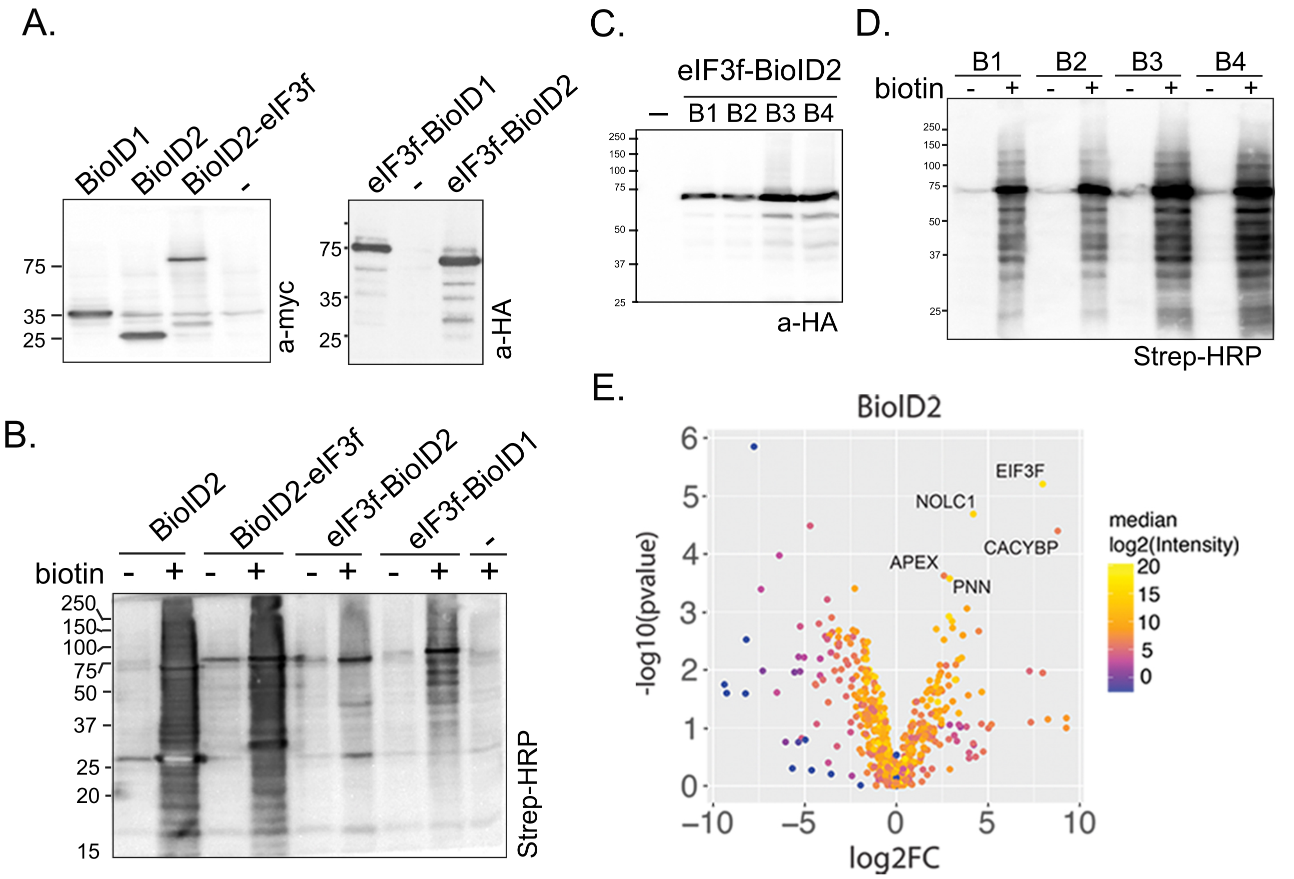


**Supplementary Figure 2: BioID1 versus BioID2 C-terminal fusion chimera of eIF3f.**

A. Transient expression in HEK293 cells of BioID1 and BioID2 fusion to the eIF3f protein (at C-ter or N-ter of eIF3f). Lysates obtained from triplicate transfections were separated on SDS gel and exogenously expressed protein detected with indicated antibody. B. Cellular distribution of the different eIF3f-BioID1 ligase constructs expressed in A. C2C12 cells were transfected with the indicated constructs. N-terminally or C-terminally fused yellow fluorescent protein (YFP) to eIF3f protein was used as control C. Western blot detection of exogenously expressed eIF3f-BioID2 (B1 to B4) from HEK293 cell lysate used for MS/MS coupled Streptavidin-dynabeads pull down. D. Streptavidin coupled HRP detection of biotin incorporation in lysates from HEK293 overexpressing BioID2 and cultured 24hrs in presence or not (+/-) of 50µM biotin. The same lysates were further used in E. We used the same protocol as previously described in figure 1. E. Volcano plot distribution of the proteins identified by LC/MS upon Streptavidin-dynabeads purification of biotinylated proteins from transfected HEK cells (supplementary figure 1A) as described in A.

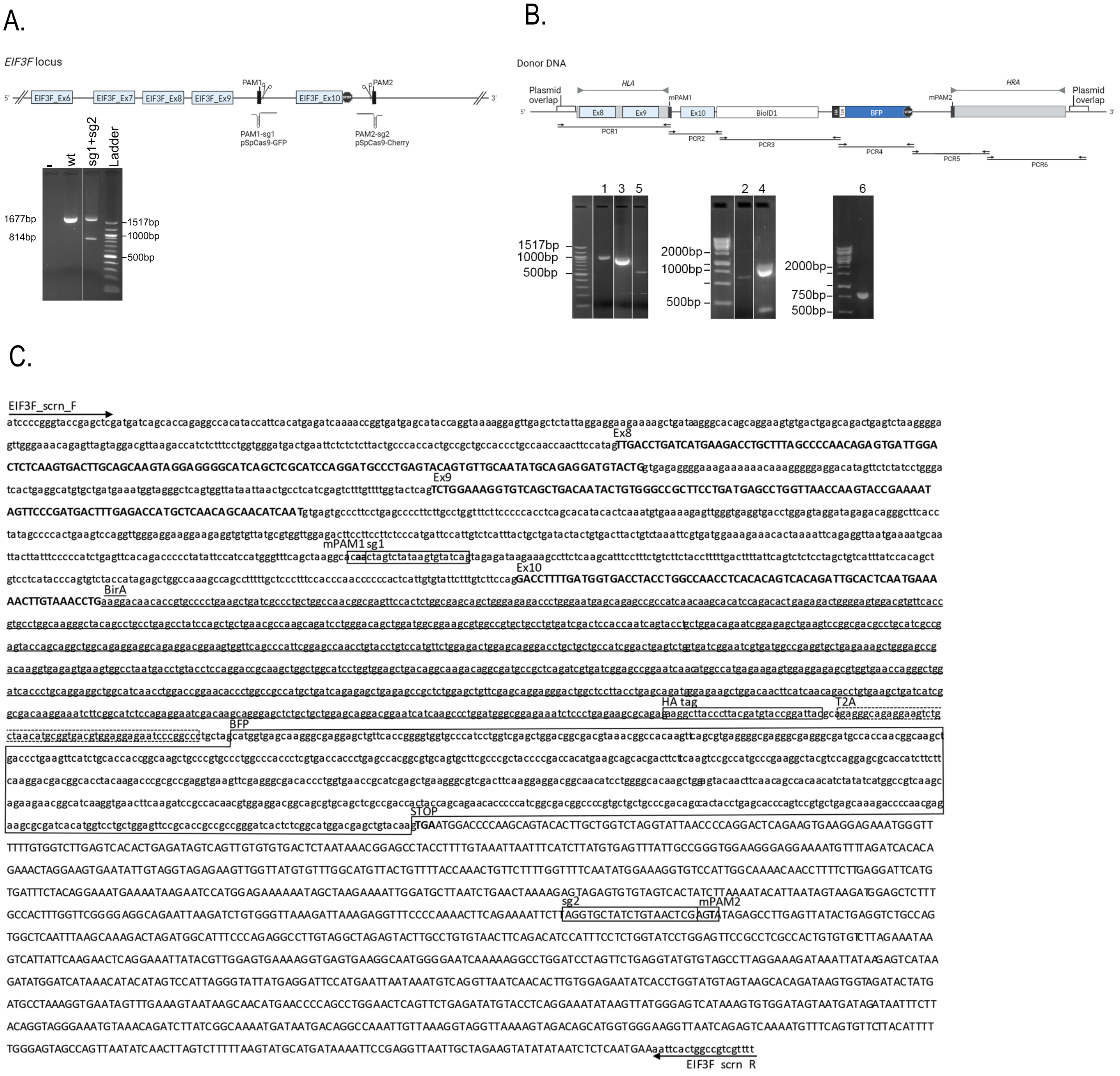


**Supplementary Figure 3: Cloning strategy and sequencing of EIF3F-BioID1 chimera.**

A. Schematic representation of EIF3F gene 3'end displaying the recombination sites for PAM1-sg1 and PAM2-sg2 on both 5' and 3' ends of Exon10. B. The Gibson cloning strategy was used to assemble the 6 different PCR amplicons of donor DNA in pUC.19 plasmid backbone. The amplified fragments corresponding to PCR1 to 6 were resolved on agarose gel before purification and assembly. C. Sequencing of the integrated construct spanning from EIF3F exon8 to 3'UTR. EIF3F_scrn_F: EIF3F screening forward; EIF3F_scrn_R: EIF3F screening reverse; Ex8: exon8; Ex9: exon9; mPAM: mutated protospacer-adjacent motif; sg: single guide RNA; BFP: blue fluorescent protein; T2A: self-cleaving 2A peptide.

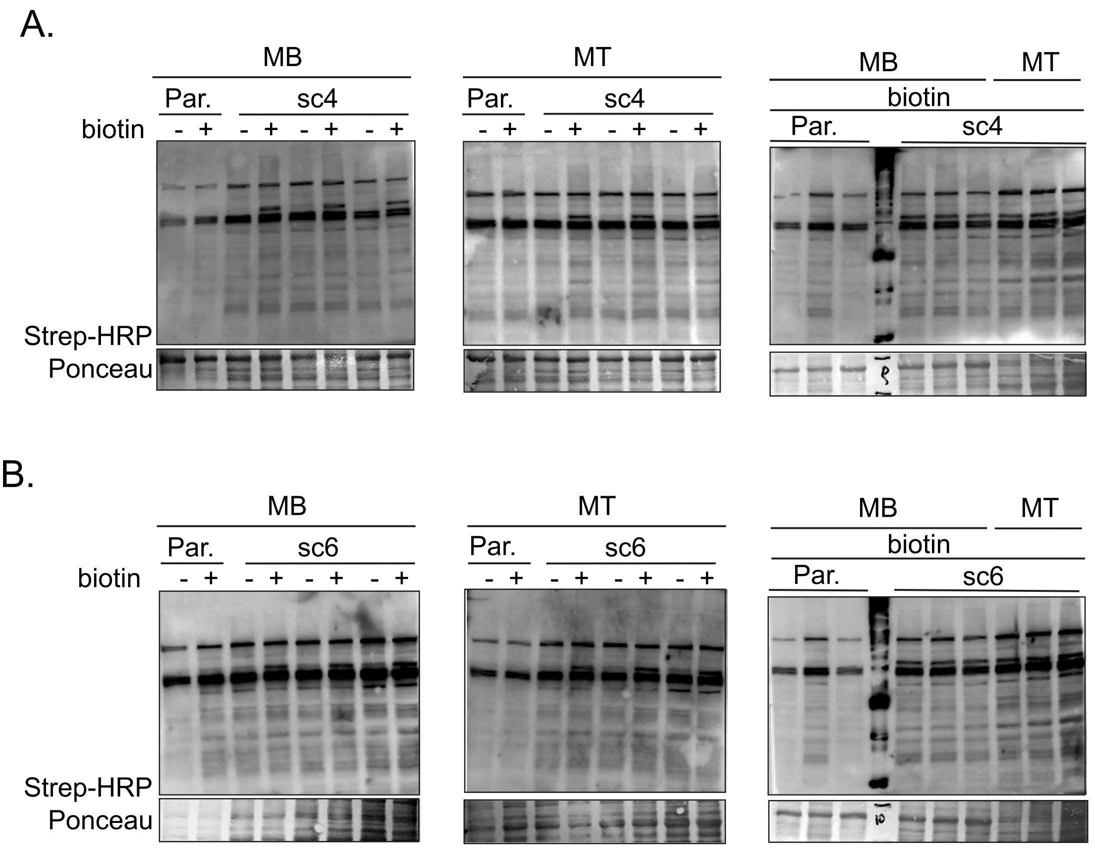


**Supplementary Figure 4: Biotin incorporation in samples used for Streptavidin-dynabeads purification of eIF3f interactors.**

A. Streptavidin coupled HRP detection of biotin incorporation form parental (Par.) and eIF3f-BioID1 knock-in sc4 lysates of MB or MT treated for 24hrs with 50µM biotin. The same lysates were further used for LC/MS coupled Streptavidin-dynabeads pull down experiment. B. The same procedure as describe in A was conducted for sc6 lysate.

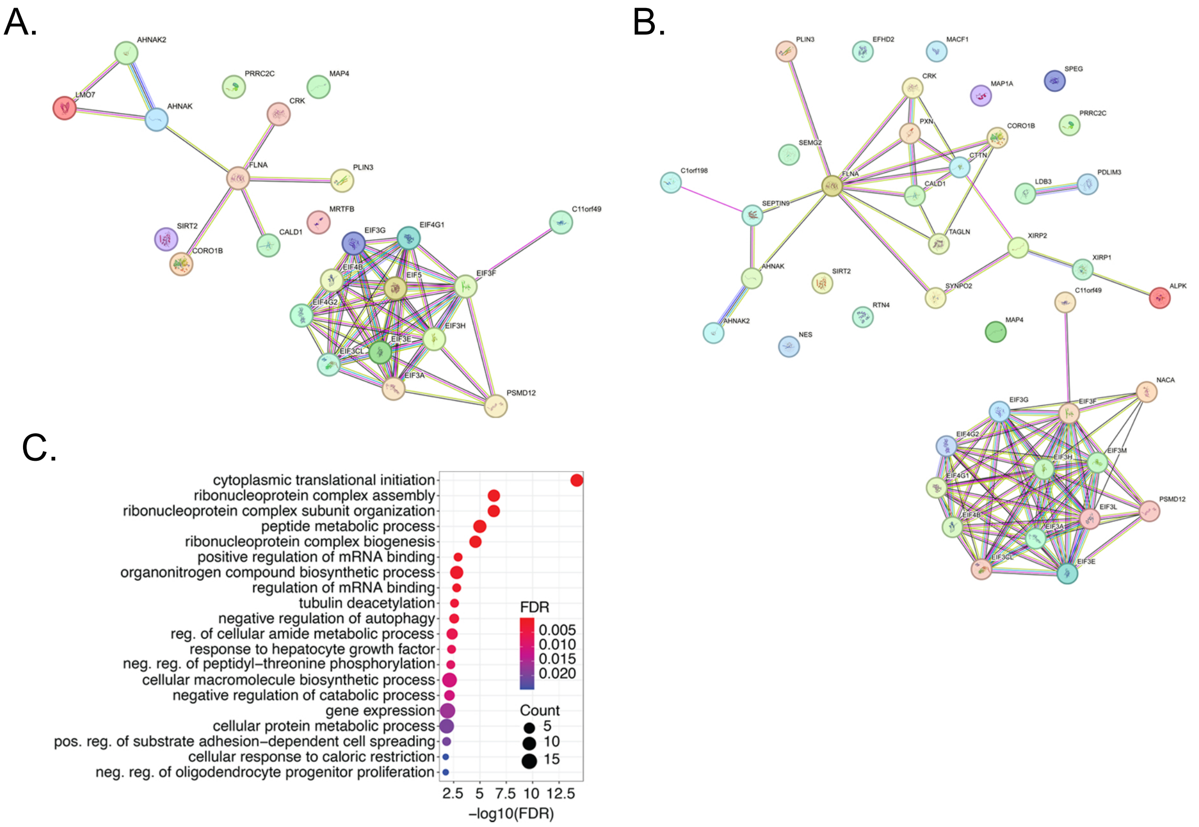


**Supplementary Figure 5: Interaction map and functional enrichment of the eIF3f interactors identified in MB and MT.**

A. Interaction map of LC/MS significantly identified (p<0.01 and log2FC>0) proteins from streptavidin-dynabeads pull-down experiment from MB (sc4/6) cultured in presence of biotin versus parental cells. B. Same as A obtained from MT (sc4/6) cells. C. Functional enrichment analysis (p<0.01 and log2FC>0) of the 21 conserved proteins identified as eIF3f binding partners in MB and MT (shown in figure 3F and 3G).

Supplementary Figure 6

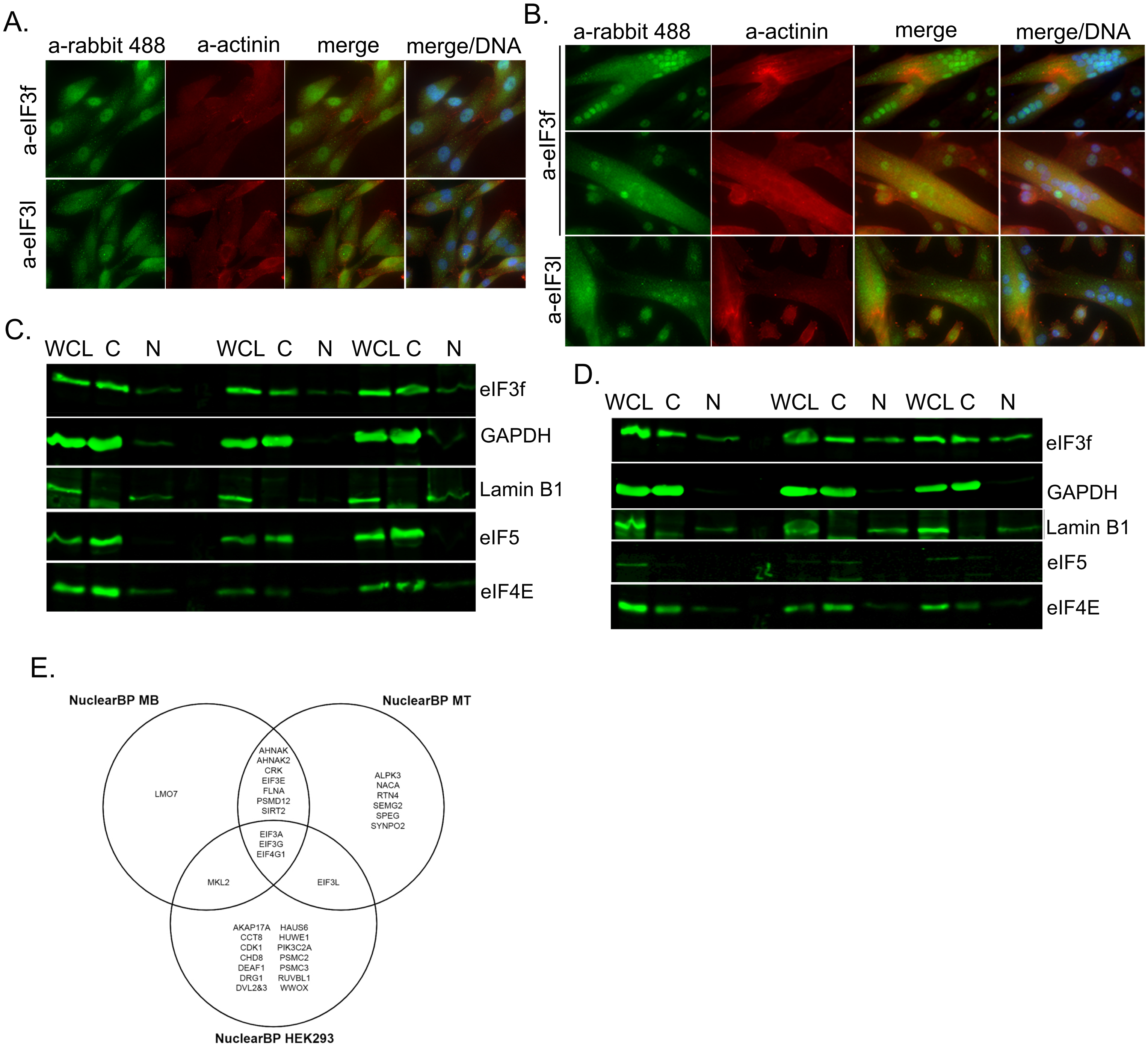


**Supplementary Figure 6: Subcellular distribution of eIF3f protein and selected identified binding partners in MB and MT.**

A. Immunodetection of eIF3 subunit F, and L (green) in parental myoblasts. Anti sarcomeric alpha-actinin-3 was visualised with Cy5 coupled anti mouse antibody and DNA was stained with Hoechst. B. The same immunodetection was conducted in myotubes. C. Whole cell lysate (WCL) or fractionated (C: cytoplasm and N: nuclear) lysates from parental MB were resolved on SDS/PAGE and probed with indicated antibodies. GAPDH and Lamin B1 were used respectively as cytoplasmic or nuclear markers. Signals from infrared (700nm) coupled secondary antibody were acquired with Odyssey (Li-Cor). D. The same experiment as in C was conducted in sc4 MT. E. Venn diagram of the nuclear binding partners (BP), according to https://www.genecards.org/ nomenclature, identified for eIF3f-BioID1 chimaera in myoblasts, myotubes and HEK293 biotinylation experiments.

Supplementary Figure 7

A.

| Protein name | log2ratio_DSB | pValue_DSB |
|---|---|---|
| sp\|P11279\|LAMP1_HUMAN | 5.46 | 9.16E-04 |
| sp\|O75112\|LDB3_HUMAN | 5.31 | 1.58E-03 |
| sp\|A4UGR9\|XIRP2_HUMAN | 5.2 | 4.84E-04 |
| sp\|P11055\|MYH3_HUMAN | 4.8 | 1.88E-03 |
| sp\|Q15772\|SPEG_HUMAN | 4.32 | 8.76E-04 |
| sp\|O00763\|ACACB_HUMAN | 3.92 | 8.34E-04 |
| sp\|Q702N8\|XIRP1_HUMAN | 3.86 | 6.95E-05 |
| sp\|P07686\|HEXB_HUMAN | 3.71 | 6.89E-04 |
| sp\|E9PAV3\|NACAM_HUMAN | 3.55 | 3.68E-04 |
| sp\|Q15050\|RRS1_HUMAN | 3.54 | 1.23E-03 |
| sp\|Q8WZ42\|TITIN_HUMAN | 3.54 | 7.88E-04 |
| sp\|Q9NQC3\|RTN4_HUMAN | 3.3 | 1.03E-03 |
| sp\|Q9NP64\|NO40_HUMAN | 3.2 | 3.39E-03 |
| sp\|Q9Y262\|EIF3L_HUMAN | 3.17 | 5.46E-04 |
| sp\|P13535\|MYH8_HUMAN | 2.98 | 2.88E-03 |
| sp\|P11182\|ODB2_HUMAN | 2.53 | 2.57E-03 |
| sp\|Q15361\|TTF1_HUMAN | 2.48 | 3.51E-03 |
| sp\|Q1ED39\|KNOP1_HUMAN | 2.17 | 2.87E-03 |
| sp\|P09651\|ROA1_HUMAN | -1.73 | 3.55E-03 |
| sp\|Q9BR76\|COR1B_HUMAN | -2.21 | 2.43E-03 |
| sp\|P16402\|H13_HUMAN | -2.57 | 1.68E-03 |
| sp\|Q9H6J7\|CK049_HUMAN | -2.69 | 8.59E-04 |
| sp\|Q5QNW6\|H2B2F_HUMAN | -3.07 | 3.18E-03 |
| sp\|P0CJ85\|DU4L2_HUMAN | -5.08 | 1.04E-03 |
| sp\|P62910\|RL32_HUMAN | -7.77 | 1.42E-03 |

B.

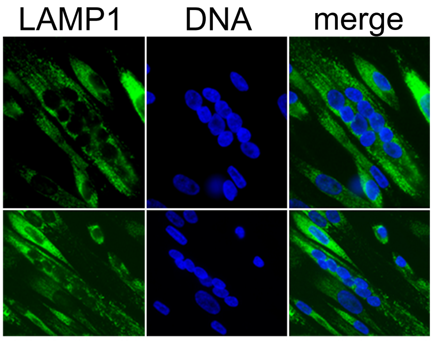


**Supplementary Figure 7: “Core eIF3f-eIF3” interactors identified in MT and MB.**

A. Table: list of interactors of the “core eIF3f-eIF3” identified in MT vs MB. Only proteins identified with high level of confidence are shown (min. med. int.>20; pVal<0.01). B. Immuno-detection of LAMP1 visualised with anti-rabbit Cy3 secondary antibody in parental differentiated (day 6) myotubes. DNA was stained with Hoechst.
